# Glycans modulate the adsorption of RBD Glycoproteins on polarizable surfaces

**DOI:** 10.1101/2025.09.30.679433

**Authors:** Antonio M. Bosch-Fernández, Willy Menacho, Rubén Pérez, Horacio V. Guzman

## Abstract

The complex interplay between glycans and protein conformational dynamics during adsorption onto polarizable surfaces opens several routes to exploring the glycans potential as molecular interactions modulators. Molecular simulations are able to dissect the interactions of Receptor Binding Domain (RBD) glycoproteins for different SARS-CoV-2 variants of concern (VoC), in both open and closed conformations, with polarizable planar interfaces. Advanced analysis projected on 2D revealed distinct adsorption mechanisms depending on the initial loci of the glycan within the protein wall. Hydrophobic surfaces facilitated stable adsorption for both RBD conformations. Conversely, hydrophilic surfaces exhibited reduced adsorption, particularly for the closed-RBD, where glycans predominantly formed hydrogen bonds. Glycans significantly modulated closed-RBD adsorption, either enhancing it by permanent tethering or impeding it depending on the two initial conformations and protein mutations (omicron). Results for the individual RBDs are shown to be consistent with simulations for the complete S1 spike glycoprotein. Our findings unveil novel glycan-mediated adsorption phenomena and provide fundamental insights into glycoprotein-surface interactions, paving the way for understanding glycan roles in protein aggregation and recognition at polarizable biological interfaces.

## Introduction

Understanding the interactions between viral proteins and their glycosylated sites have recently driven significant efforts by the molecular simulation community in comprehending novel intrinsic functions of glycans.^1–5^ Those interactions are highly dynamic and environment dependent, as they have an amphiphatic nature that allows glycans to interact with polar and non-polar protein residues or surfaces.^4,6,7^ Particularly poorly understood remains the question of how glycans modify the adsorption of glycoproteins onto polarizable surfaces.^8^ Dissecting the interactions of glycans in the environment provided by hydrophobic and hydrophilic surfaces is crucial to uncover the polysaccharide-chain ability to enhance or weaken the adsorption process. A critical challenge in studying adsorption processes is defining proper initial conformations of the most flexible molecules (glycans) in the glycoprotein-surface system.^8,9^ Similar questions have been addressed to model the glycoprotein interaction in bulk water,^3,9–11^ with receptor proteins^12–14^ and related setups.^8,15^ Each of those studies revealed important aspects of glycan interactions, as for example the shielding mechanism of glycans towards the immune system defenses which remarked the critical role of glycans during the vaccine development period against COVID-19. ^1^

These findings have provided the momentum for addressing novel questions in glycosylation from the computational biophysics perspective. Among them is the effect of combining a commonly semi-flexible molecule (proteins)^16,17^ with highly flexible ones (glycans).^18^ The challenge here is how to evaluate the effects of flexibility on the interaction in 3D bulk. Polymer adsorption theory and simulations^19–21^ provide us tools to evaluate this phenomenology by tackling the interactions onto planar surfaces. This analysis benefits from a simplified representations that enable us to guide the structural analysis of the adsorption configurations of biopolymers (glycoproteins, nucleic acids and membranes). Model polarizable surfaces^22^ offer an effective way to address the effect of hydrophobicity/hydrophilicity. They have been used in the characterization of the adsorption of biopolymers onto planar surfaces and membranes,^21,23^ to tackle the electrostatic interactions of the VoC spikes with planar and charged surfaces,^24^ and in our recent study of the adsorption of SARS-CoV-2 Receptor Binding Domains (RBD) for the WT, Delta and Omicron variants of concern (VoC) in an open conformation. ^8^ Recent experimental results with High-speed AFM^25^ characterized the interaction between the viral trimeric spike protein S1 and inanimate surfaces, which revealed the highly dynamic transitions between open and closed conformations close to a substrate. The highly dynamic interaction of the RBDs onto mica was previously observed by Hinterdorfer and co-workers for different VoCs, and also quantified the spike and host angiotensin-converting enzyme 2 (ACE2) interactions.^11^

These experimental findings motivate the present study, where we employ molecular dynamics (MD) simulations to examine the interaction of individual SARS-CoV-2 RBD glycoproteins in the closed conformation and the whole S1 spike glycoprotein with model polarizable surfaces (see Figure 1). Whereby, we previously gain insights from MD simulations of the open conformation. ^8^ Our approach builds up on the understanding of the adsorption of several single RBD conformations and the piecewise reconstruction of the same process for the whole S1 spike glycoprotein. This strategy allows us a comprehensive understanding of the S1 adsorption and, more importantly, to explore the interplay of the protein-glycan and glycan-glycan interactions together with the direct protein- and glycan-surface interactions in shaping the process at both RBD and S1 levels. To this end, we employ basic^23,26^ and develop novel 2D analysis tools to quantify the morphological changes and the number, nature and spatial distribution of the contacts formed as a function of the initial location of the glycan (in the RBD head or the Receptor Binding Motif –RBM– for the closed-RBD, see Figure 1C) for each polarizable surface and RBD VoCs.

**Figure 1.**
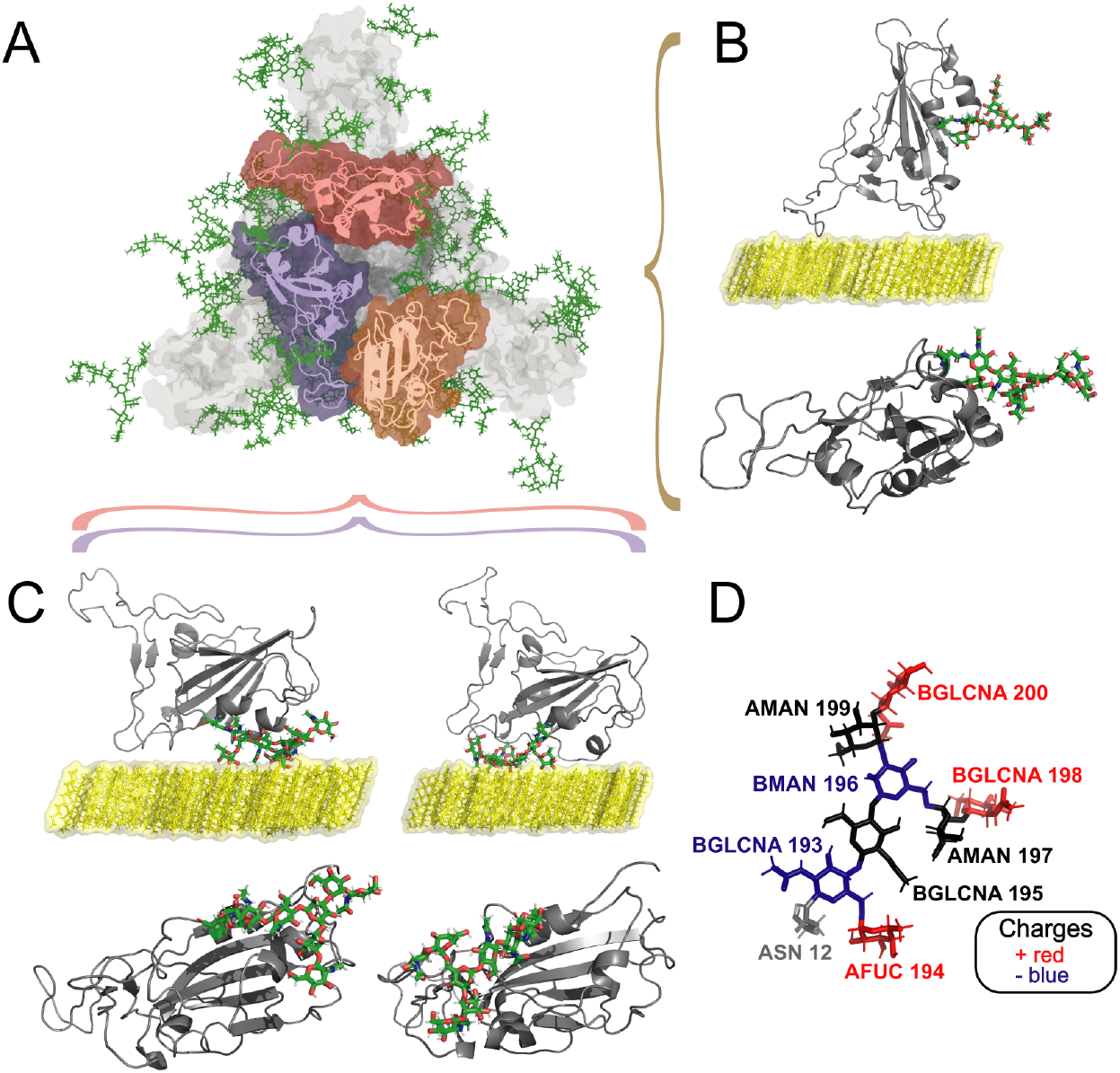
Protein–glycan complexes simulated in this study in the presence of polarizable bilayers (PBLs). (A) Bottom view of the S1 Spike glycoprotein (S1S), indicating with different colors the three RBD monomers, where one RBD is in its open configuration (orange), and the other two are in closed configuration (red and purple). The non-RBD regions of the S1S are in gray, and glycans are in green. (B) Front and bottom view of the isolated open-receptor binding domain (RBD), previously analyzed in detail in our previous work.^8^ (C) Side and bottom view representations of the RBD in closed configuration, showing the glycan positioned at the RBD head (left) and the RBM (right), respectively. Glycan colors in (B,C) follow an atom type format, where green corresponds to carbons, red to oxygen, and white to hydrogen atoms. (D) Detailed view of the RBD’s glycan, including residue names and positions labeled; blue, red, and black colors are used for negatively charged, positively charged, and neutrally charged glycan residues, respectively. The amino acid linking the glycan to the RBD is shown in gray.

Our main result is the identification of a dual glycan capacity, to act either as a glue that dominates the whole glycoprotein adsorption strength during intermittent glycan-surface tethering patterns, or as a blocking element preventing the protein-surface interaction. While glycans do not significantly influence the adsorption of open-RBDs due to its location far from the protein-surface contact area,^8,27^ they are crucial for tuning the adsorption in the closed-RBD conformations. Specifically, our analysis reveals stable adsorption to hydrophobic surfaces of both open- and closed-RBD glycoproteins. While the flexibility of the receptor binding motif in the open configuration does enhance the adsorption strength, ^8,28^ the contact region of the closed-RBD shows no significant structural changes. Hydrophilic surfaces are less prone to adsorption, specially in the closed conformation of the glycoprotein. Here, the glycan location plays a key role due to two different effects: its monosaccharides form the majority of hydrogen bonds and they can block the interaction of the surface with hydrophilic areas of the RBD. The strategy of using planar interfaces to reduce the degrees of freedom of the short-range interactions in glycoproteins allow us to show the glycans’ high flexibility, to track in detail its interaction with the protein, other glycans and the surface, and to reveal how these effects combine to drive or impede the adsorption.

The novel adsorption mechanisms identified between glycoprotein and surfaces, together with the very recent development of methods that allow tracking glycans with angstrom resolution,^29^ provide the starting point to understand these processes in heterogeneous interfaces where glycans could induce protein aggregation, multi-protein assemblies, and misrecognition of protein-based drugs.

## Results

### Morphological changes and contact formation during adsorption

The simulated RBD glycoproteins along this work consider three VoCs and three main conformations of the RBD, namely, open, closed-head and closed-RBM (see Fig. 1). In Fig. 1D, we have also displayed a close-up view of the N343 glycan^5^ residues and polarities. We produced trajectories for each system (see Methods for details) and used them to extract representative morphological changes and detailed information about the contacts formed during the adsorption process for both hydrophilic and hydrophobic surfaces.

The morphological analysis is based on the parallel and perpendicular radii-of-gyration, *R*_*g*∥_, *R*_*g*⊥_, calculated for trajectories based on 3 replicas and shown as individual dots, together with their time averages, in Fig. 2). Figs. 2 A,C and B,D correspond to the two different initial positions of the glycan in the RBD glycoprotein in closed conformation, namely, the glycan is located at the head position (A,C) or at the RBM site (B,D), as previously described in Fig. 1C. In terms of the ratio 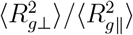 (deformation) for the proteinaceous component of the glycoproteins, the closed RBD conformations (white symbols) are clearly less flexible than the open ones (gray symbols): the relative percentage of the proportion of deformation ratios between closed-head and open is in the range ≈ 15-25% (depending on the VoC) while for closed-RBM and open is ≈35-45% (see SI Table S1).

**Figure 2.**
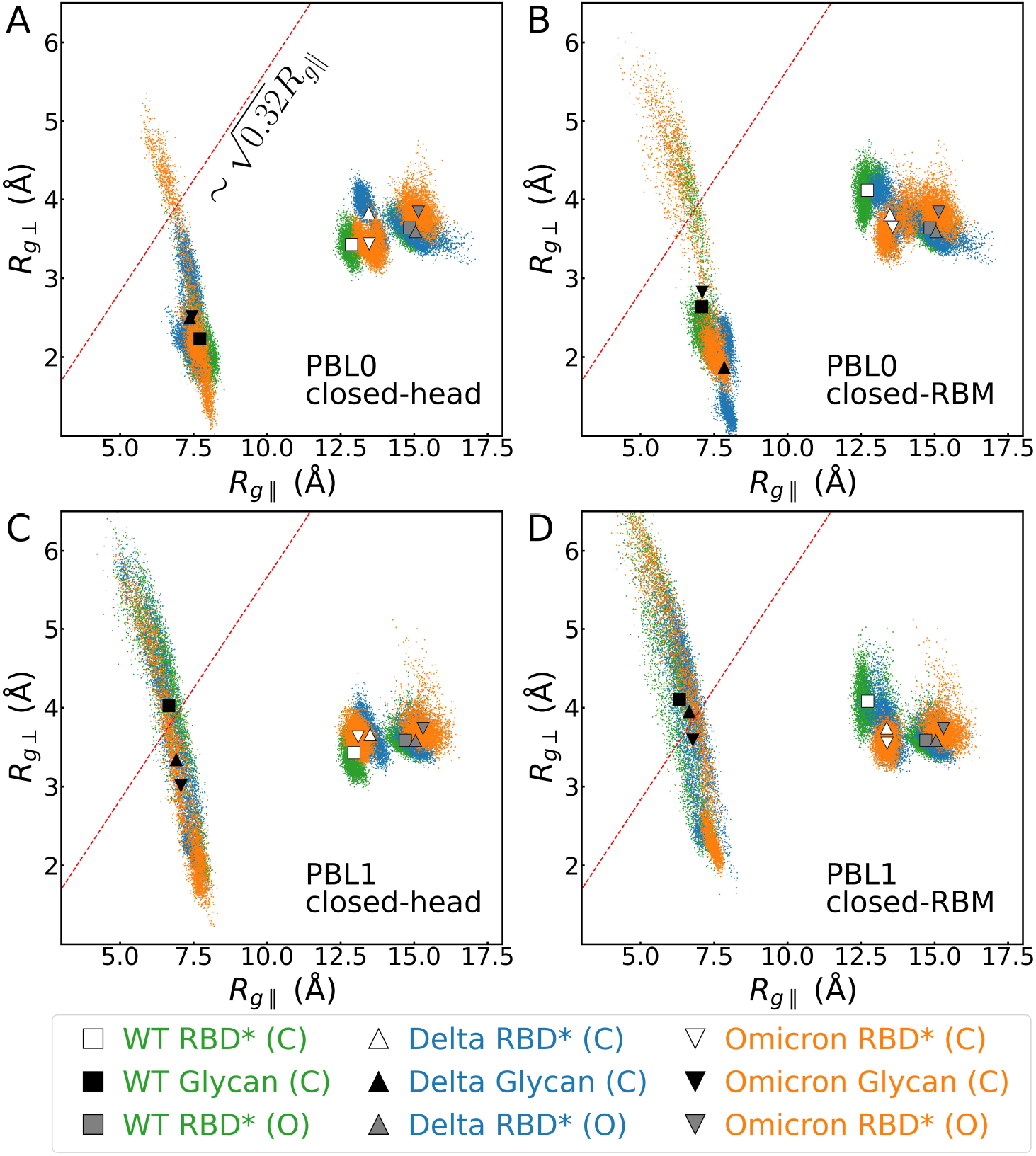
Parallel and perpendicular radii of gyration of the RBD protein in its open configuration in the absence of glycans (grey symbols), in its closed configurations (white symbols), and the glycan alone (black symbols) of the simulation with closed-RBD. Small green, blue, and orange dots correspond to the data at each frame, and the symbols represent the mean values for WT (square), Delta (triangle), and Omicron (inverted triangle) variants, respectively. Red lines indicate the limit 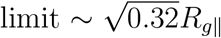 for the semiflexible polymer regime. Panels show closed-RBD with the glycan between (A) a hydrophobic surface and the RBD’s head, (B) a hydrophobic surface and the RBD’s RBM, (C) a hydrophilic surface and the RBD’s head, and (D) a hydrophilic surface and the RBD’s RBM. Note that the results of radii of gyration of the proteins do not consider the whole protein, but only the contact regions with the surface (cases marked with * in the legend).

Considering the surface polarizability at the hydrophobic interface, as reported previously for the open conformation, ^8^ the adsorption is stronger than in the hydrophilic case. Note that in this work, we also considered the glycan-surface interaction (black symbols and corresponding distribution). *R*_*g*∥_ and *R*_*g*⊥_ along the trajectory (see Figs. 2) show much higher fluctuations for the glycan compared to the protein, evidencing the highly flexible nature of the glycans. However, the average values (black symbols for glycan, white and gray for the protein in Figs. 2 A and B) remain in the semi-flexible regime. This result can be understood from the fact that, in the great majority of these trajectories, the glycan is located between the protein and the surface. There are very subtle differences in the ratio 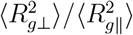 between the two glycan locations, namely, a slightly more flattened protein when the glycan is at the head (Fig. 2A), resulting from the fact that RBM region is more flexible and spreads more than the head one (See also SI Fig. S1). On the contrary, in the closed-RBM conformation (Fig. 2B) the glycan prevents the direct interaction between the RBM region and the surface and, hence, its deformation during adsorption is around 1.5 times lower for both WT and omicron (see SI Table S1).

For the hydrophilic surface, we observe a significantly different behavior in the glycan adsorption for the closed conformations: The three VoCs present a huge fluctuation pattern, with the WT variant lying outside the boundary of semiflexible adsorption, while delta and omicron variants are very close (Figs. 2C and D). The glycan fluctuations can be associated with tethering conformations where the glycan interacts with the surface by forming hydrogen bonds, and which are highly dynamic. Focusing now on the protein adsorption, there are clear differences between the two glycan locations for the WT: the ratio 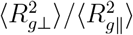 increases 1.48 times (see SI Table S1) when comparing the head and RBM locations, reflecting the fact that the head-contact region is more hydrophobic than the RBM one. These differences are less pronounced in the omicron variant, mainly due to the the mutations bringing charged residues and intermittent formation of H-bonds.

Fig. 3, compares both the total number of contacts and the contact area (see Methods) of the RBDs in 3 initial conformations, open, closed-head and closed-RBM interacting with two model surfaces. Divergent from the open RBD conformation, ^8,27^ the N343 glycan in the closed conformations plays a crucial role in promoting or blocking protein adsorption.

**Figure 3.**
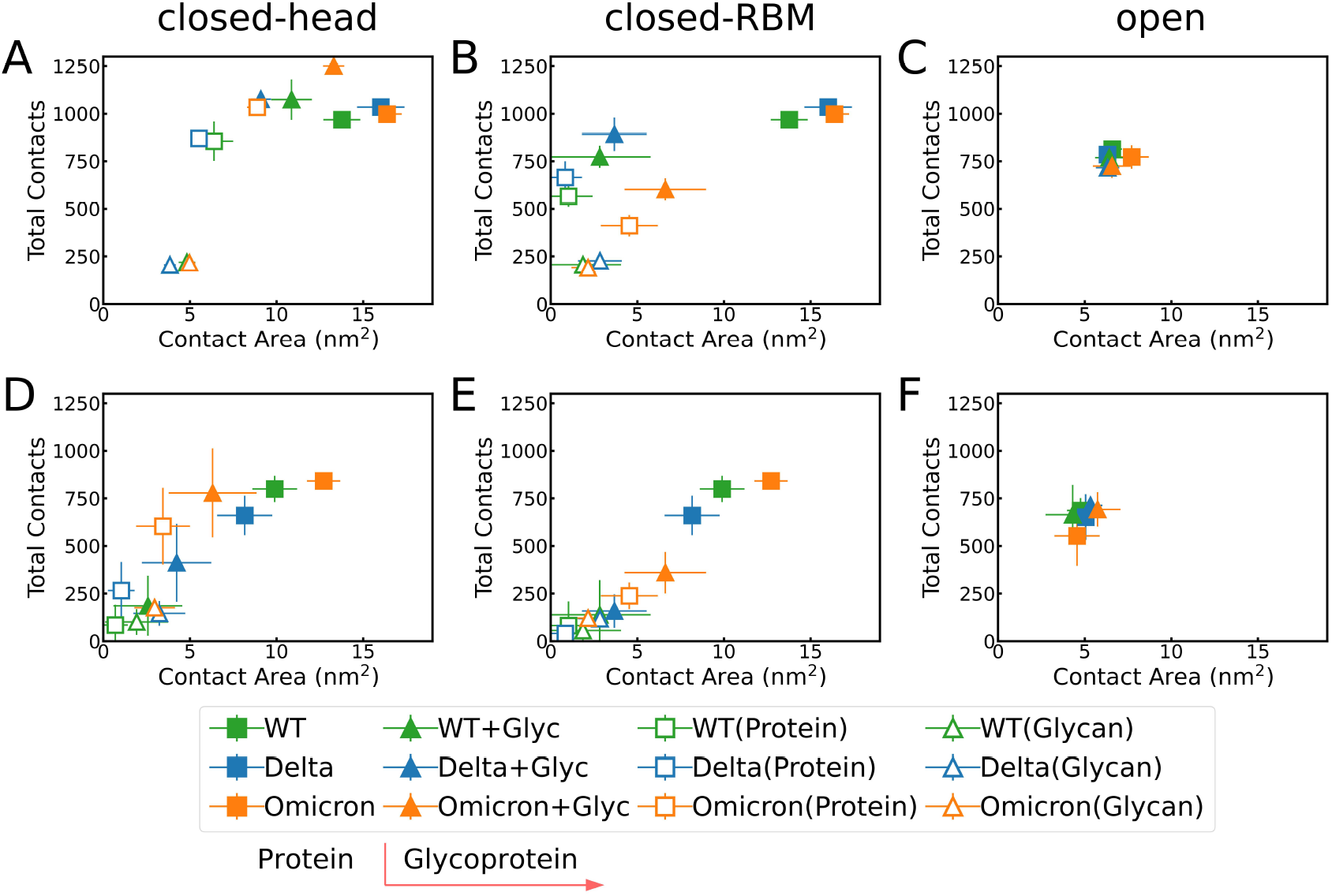
Total number of contacts versus contact area for interactions with (A–C) hydrophobic surfaces and (D–F) hydrophilic surfaces. Solid squares represent data from simulations without glycans, while solid triangles correspond to simulations including glycans. Empty squares and empty triangles indicate the contributions of the protein and the glycan, respectively, in the protein–glycan complex simulations. Panels show results for the RBD in (A, D) the open configuration, and in the closed configuration with the glycan positioned at the (B, E) surface–head RBD interface, and (C, F) surface–RBM RBD interface.

Focusing on the two close conformations for the hydrophobic case, the closed-head one doubles the contact area of the closed-RBM conformation (full triangle symbols denote the glycoprotein in Fig. 3). However, the total number of contacts decreases from ∼20% (WT and Delta) to ≈50% (Omicron) when the glycan is initially located at the RBM region. As expected, this difference is related to the mutations in omicron’s RBM region, which turn it more hydrophilic (see SI Tables S3 and S4). When the glycan flattens between the protein and the surface, it forms hydrophilic interactions with the protein, preventing the protein adsorption on the surface. This conclusion is confirmed by the separated analysis of glycan and protein (empty symbols in Fig. 3) that shows for closed-RBM a significant reduction in the protein contacts and in its contact area. These changes are drastic for WT and Delta (reducing them to minimal values) while, in the case of Omicron, the protein contact area is only half the one for the close-head case.

Both the total number of contacts and the contact areas reduce for all VoCs at the hydrophilic surface (compared to the hydrophobic one), as shown in Figs. 3D, E and F. Comparing the two glycan locations, both quantities decrease when moving from the closehead to closed-RBM conformations, but the reduction in contact area is not drastic due to the fact that the hydrophilic interactions (associated with the formation of hydrogen bonds) are dominated by the glycan, as discussed below.

The footprints or 2D-densities of the top 5 residues –the location of their center-of- mass (COM) along the trajectory–, for WT and omicron forming more contacts with the hydrophobic surface, together with the position of the glycan’s COM, are shown in Fig. 4. The data extracted from the trajectories for each system (closed-head, closed-RBM, and open, as specified in Table S11) reveal the significant dependence on the conformation and the initial glycan location. The insets in Fig. 4 display the time dependent distances with the surface of these residues (blue) and the glycan (pastel-red curve).

**Figure 4.**
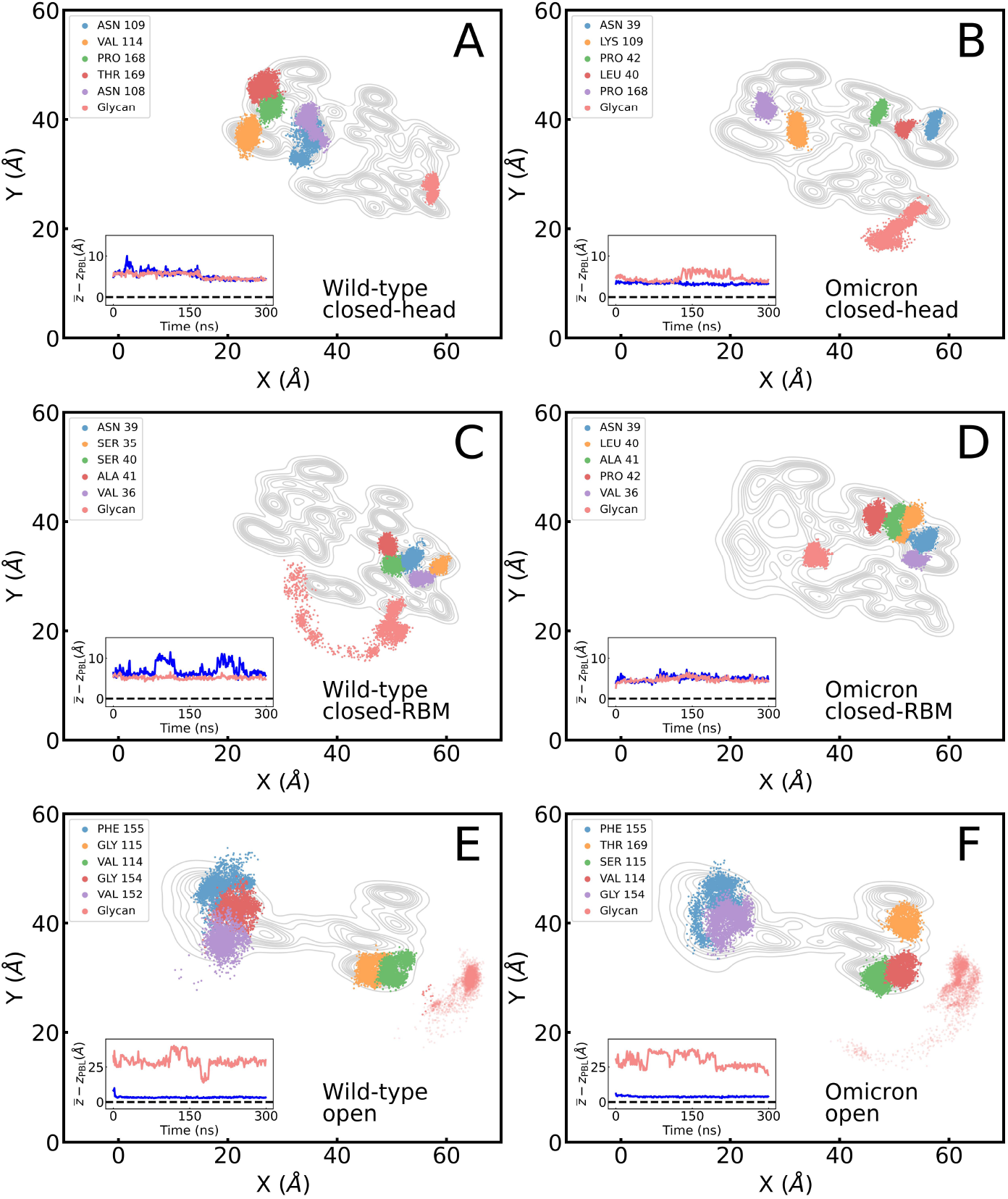
Footprint of the top five protein residues with the highest number of contacts to the hydrophobic surface and center-of-mass (COM) of the glycan for the closed- and open-RBD configurations for WT (left column) and Omicron variants (right column). Insets show the average distance of these residues (blue) and the glycan (red) to the surface. Panels display results for (A, B) closed-RBD with the glycan between the RBD head and the surface, (C, D) closed-RBD with the glycan between the RBM and the surface, and (E, F) open-RBD configuration. Contours of the contact regions of closed and open RBDs are shown as a reference map for the protein.

In the closed conformations, both distances go mostly along with each other (see insets in Figs. 4A, B, C, and D). This behavior is completely different from the one in the open conformations (Figs 4E and F), tackled in our previous work,^8^ where the glycan is attached on the side (Fig. 1B): The size of the RBD’s side-wall is commensurate with the length of the modeled polysaccharide chain, precluding a direct interaction between Glycan and surface for open conformations. The situation is different for the closed conformations, where the glycan is initially located between the protein and the surface. Here, depending on the glycan initial loci and the VoC, we observe a diverse phenomenology. For the close-head configuration (Fig. 4A), the 5-top interacting residues are located at the RBM for WT, while, for omicron, 3 of them are found at the head-side. This distinct behavior results from the combined influence of mutations at the head site, which turned more hydrophobic (see SI Table S3), and the glycan mobility. The glycan-surface distance (pastel-red curve inset Fig. 4B) illustrates the process where the glycan squeezes out from its initial location towards the liquid-surface interface and jumps out-of-contact for over one third of the trajectory.

A quite different behavior is found in the closed-RBM case (Figs. 4C and D). Here, the glycan flexibility contributes to hold the WT glycoprotein in contact with the surface during different unbinding events (Fig. 4C). The protagonists of those events were 2 serines (ResIDs 35 and 40) located at the WT’s head. In omicron, two mutations (LEU 40 and PRO 42) enter the top-5 contacts and contribute, with their non-polar character, to provide a stable adsorption during the whole trajectory (see inset of Fig. 4D). Representative snapshots of the adsorption behavior for the hydrophobic surface are illustrated in the SI Fig. S4.

The results for the hydrophilic surface are shown in SI Fig. S2. Here the adsorption behavior follows, in general, an oscillatory pattern as the one previously reported for the open conformation.^8^ However, in the close conformations, the glycan plays an active role, trying to keep the glycoprotein in contact with the surface (see the snapshots in SI Fig. S5). As can be expected from the almost identical sequences in the contact region (see SI Table S3 and S4), the adsorption behavior of delta (SI Fig. S3 and snapshots in SI Fig. S5) is very similar to WT on both surfaces. To summarize this analysis, we have quantified the angular interactions of each monosaccharide of the N343 glycan (SI Fig. S6) in order to further confirm the main message: the amphiphatic behavior of this glycan onto either hydrophobic (with angles less than 45 degrees) and hydrophilic ones (with larger angles).

### Fluctuation analysis and Hydrogen bonding

The identification of the residues with highest RMSF values (Fig. 5) provides further insight into the contact analysis presented above. Correlating the number of contacts with regions with larger fluctuations illustrates whether fluctuations promote more contacts at the interface or those contacts are rather based on highly localized short-range interactions with less flexible protein regions. Fig. 5 presents, for WT and Omicron adsorbed on the hydrophobic surface, the footprints of the five protein residues with the highest RMSF values, the areas covered by them and by the glycan COMs (see legends for quantitative values) and a profile with the RMSF values for all protein residues in the contact region. The same analysis can be found in the SI for the hydrophilic surface (Fig. S7) and the delta variant (Fig. S8).

**Figure 5.**
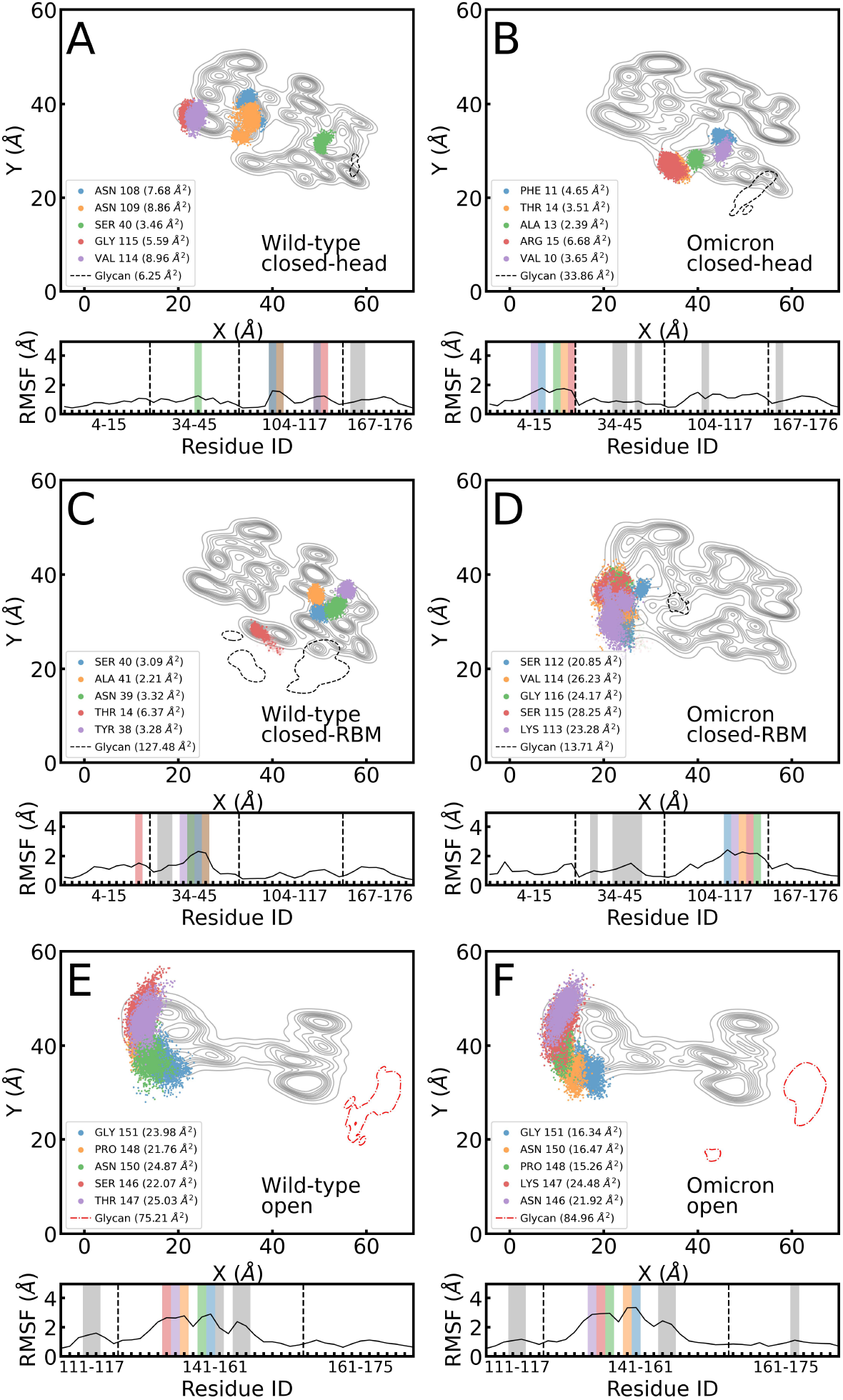
Footprint of the top five protein residues in the contact region with the highest RMSF values and area covered by glycan (dashed line) for the closed-RBD and open configurations onto the hydrophobic surface for WT (left column) and Omicron variants (right column). Panel ordering is consistent with Figure 4. RMSF values for residues in the contact region are shown below each footprint, with colored regions highlighting the residues shown in the footprint and grey regions corresponding to residues with the highest number of surface contacts (Fig. 4). Red dot-dashed lines in (E) and (F) are a reminder that the glycan in the open conformations is not adsorbed (normal distance *>* 15 Å).

According to the 2D fluctuation areas for the closed-head conformation (Figs. 5A and B), the glycan in WT fluctuates less at the interface than in omicron. The RMSF profile for the closed-head conformation in omicron shows that all of the residues with highest RMSF are located in close proximity to the area covered by the glycan. None of these residues are among the top-5 in number of contacts (Fig. 4B). This result highlights how glycans can prevent sterically the approach of residues to the surface, particularly for the 3 hydrophobic residues (VAL 10, PHE 11, and ALA 13) captured in Fig. 5B. This behavior is not found in WT (Fig. 4A), where 3 (ASN 108, ASN 109, and VAL 114) out of 5 residues with larger fluctuations at the interface are also the ones achieving more contacts.

For the WT closed-RBM conformation (Fig. 5C), glycan fluctuations are in the spotlight as its area is 2 orders of magnitude larger than the ones of the protein residues with the highest RMSFs. Here, glycan fluctuations are effectively promoting the adsorption of residues, because the glycan extends its interaction with the surface through different areas than the location of residues at the RBM. As seen before for the close-head conformation: again, 3 (ASN 39, SER 40, and ALA 41) out of the 5 more fluctuating residues coincide with the residues with more contacts Fig. 4C). Here, the adsorption is mainly driven by the head region that includes the most hydrophobic residues, which also pull neighboring hydrophilic ones during their adsorption process. In the case of omicron, Fig. 5D illustrates the effect of neighboring residues (ResIDs: 112-116) in promoting fluctuations: Larger RMSFs are found in the vicinity of mostly hydrophilic residues at the RBM region, including the GLY-to-SER mutation of residue 115. On the contrary, interactions with the hydrophobic surface (mutations turned the head region in omicron more hydrophobic) led to significantly smaller RMSF values, around half of the top ones (gray areas in the RMSF profiles).

The open conformations (Fig. 5E and F) are, by far, the ones where protein residues fluctuate the most, with RMSF values much higher (maxima around 3 Å) than for the closed conformations (that remain close to 2 Å), with the exception of the closed-RBM omicron’s case. These results illustrate the case where the glycan, due to its location on the side of the RBD protein, plays a minor role in the RMSF footprints and are shown for comparison.

Nonetheless comparing their RMSF profiles, the open conformations for both variants reach values around 3 Å while the closed ones remain in the vicinity of 2 Å. An important remark for the glycan RMSF areas shown for the open conformation are only a projection on the surfaces place. However, the z distance is not within 15 Å from the surface and hence are shown in red dot-dashed lines (see Fig. 5E and F).

Fig. 6 shows the protein and glycan residues that preferentially form hydrogen bonds (H-bonds) with the hydrophilic surface for WT and Omicron for the three conformations described above. Circles are centered at the COMs of the residues, and their size represents the percentage of the frames along the trajectory where this residue established an H-bond. Contour lines for the corresponding RBD protein and the glycan (dashed line) provide a reference to identify the areas that dominate H-bond formation. At variance with the open conformations studied previously^8^ and represented here by a single trajectory (Figs. 6E and F), H-bond formation for the closed configurations is dominated by the N343 glycan (see Fig. 6 and the the SI Tables S6, S5 for quantitative values).

**Figure 6.**
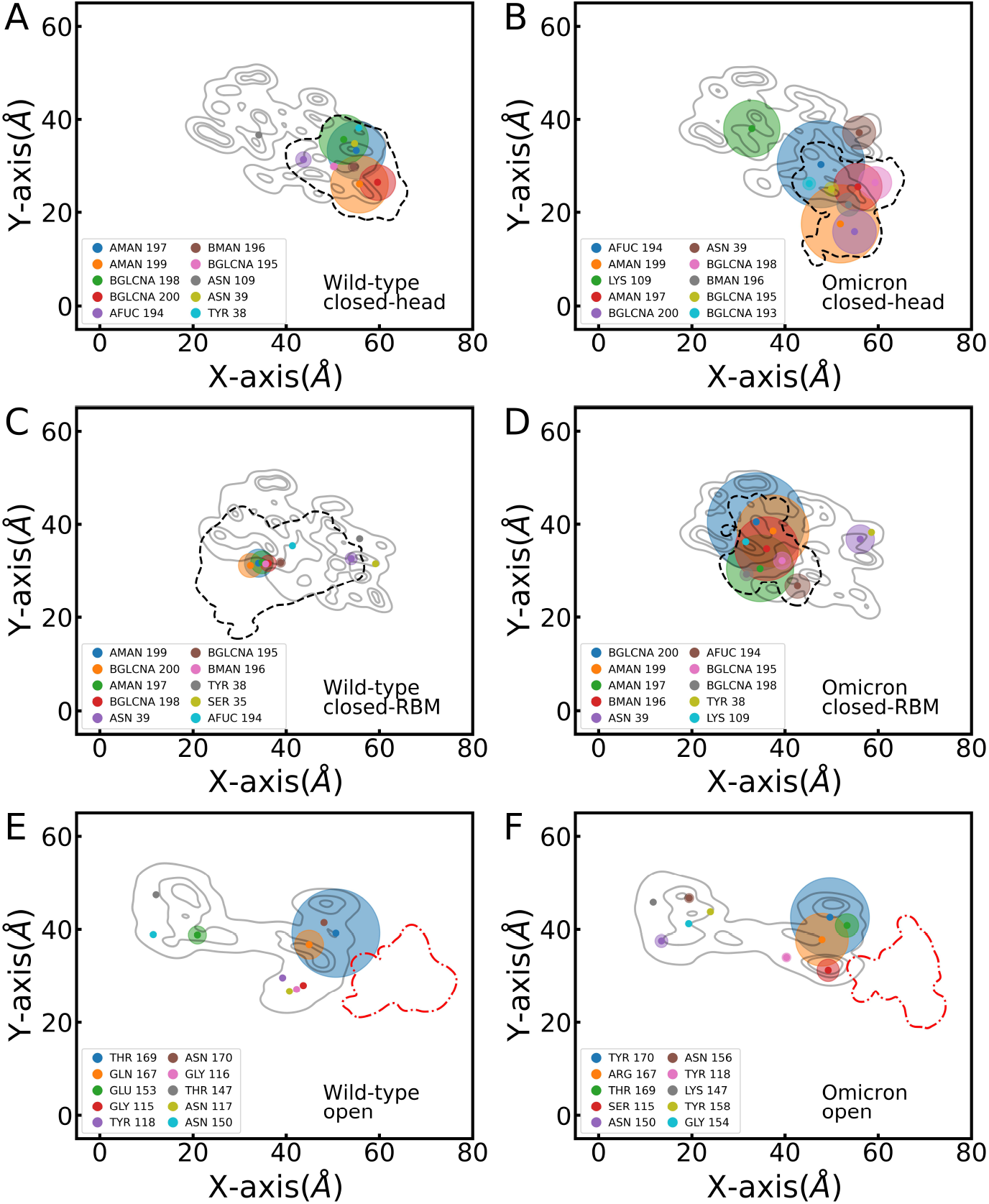
Hydrogen bonds occurrence between the protein and the surface during the simulation time (shown as %). H-bonds are represented in circles centered at the center-of-mass of each residue with proportional size to the percentage of H-bonding during the trajectory. The three contour levels in gray represent the mean positions of the protein, and the single contour in segment-black indicates the mean position of the glycan. Panel ordering is con-sistent with Figures 4 and 5, with WT in left panels and Omicron in right panels.

Starting with the head-RBD, seven glycan residues are the ones with the highest percentages in WT (Fig. 6A). Mutations from hydrophobic to hydrophilic residues at the RBM in Omicron compete with the glycan, with the LYS 109 residue appearing in third position and reaching nearly 35% of H-bonds along the trajectory (Fig. 6B). This result explains the significantly larger contact area of the omicron variant over the WT discussed above (see Fig. 3). Protein mutations clearly affect also the glycan conformation and H-bond formation, with the glycan AFUC residue reaching in the omicron case a percentage around 5 times that of WT. The glycan dominance reflects also in the H-bond distribution within the RBM region: the balance found between the two groups (left and right sides) in the open conformation (see Figs. 6E and Fig. 9 in ref. ^8^) is lost in the close conformations, where the glycan clearly defines the H-bonding rich interaction regions. Similar trends are found for the closed-RBM conformation, albeit a significant reduction in H-bond formation in WT compared to the close-head. In fact, the closed-RBM WT H-bonds occurrence clearly show the versatility of the glycan to monopolize most H-bond interactions. The study of the delta variant (SI Fig. S9 and S7) leads to the same conclusions, with a clear predominance of the glycan in H-bond formation for the closed conformations and a rather balanced H-bond occurrence for the open RBD.

### Glycan interactions with the surfaces and the protein walls

After the analysis of the adsorption of the glycoprotein as a whole, we focus our attention on the glycan and characterize, at the level of a single monosaccharide, the interaction with the surface and protein during adsorption. Fig. 7 shows, in the closed conformations of WT and Omicron, the specific contacts formed by each of the glycan residues with the hydrophobic (Figs. 7A and B) and hydrophilic (Figs. 7C and D) surfaces. General trends, like the drastic reduction in the number of contacts in the hydrophilic case irrespective of the VoC, have been already identified in the literature for proteins^8^ and surfactants,^22^ or in our own analysis for the glycoprotein above, that showed a reduction of the number of contacts for both closed conformations. Fig. 7 confirms that these trends also apply to the N343 glycan, demonstrating its amphiphatic character. Here, the in-depth analysis at the residue level provides more insight into the role of the glycan in the process. While for the hydrophobic surfaces, there are no significant variations in the number of contacts associated with each of the monosaccharides of the glycan chain, they can be clearly spotted in the hydrophilic case, particularly for the WT variant. This indicates that the glycan is not completely adsorbed, and there are intermittent interactions with the surface where the last monosaccharides of the chain (the ones located further from the anchoring point to the protein residue ASN 12) help tethering the WT-RBD (See SI Fig. S5A). This is also consistent with the high fluctuations of the parallel and perpendicular radii of gyration of the glycan in Figure 2). For omicron in closed-head conformation (Fig. 7C), the enhanced hydrophilic character of the RBM adds to the glycan contact formation with the hydrophilic surface and hence promotes adsorption (SI Figs. S5B). For the closed-RBM conformation, the head region –with its more hydrophobic character with respect to WT– is exposed, what hinders omicron adsorption on the hydrophilic surface with respect to the closed-head conformation. The analysis for the delta variant (SI Fig. S10) shows again the amphiphatic behavior of the glycan, that is further confirmed by our analysis of the angles of the sugar rings with respect to the surface normal (SI Fig. S6).

**Figure 7.**
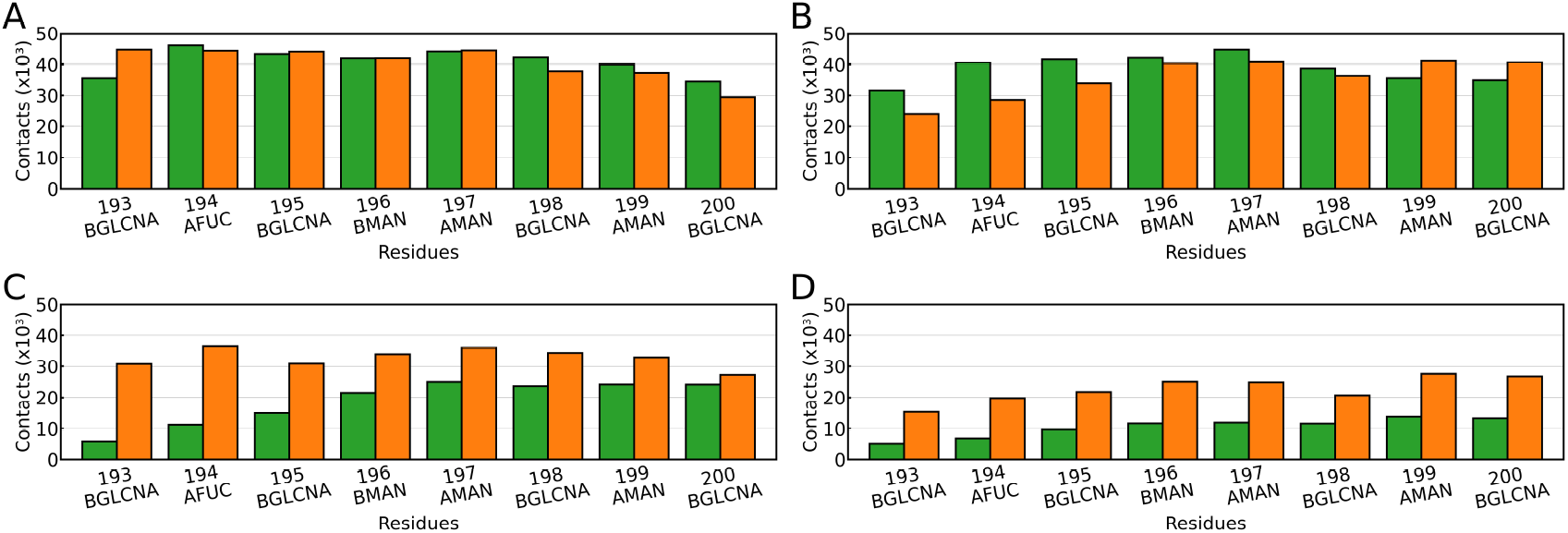
Glycan contacts with the surface when positioned between the surface and (A, C) the RBD head, or (B, D) the RBM region. Panels (A) and (B) correspond to a hydrophobic surface, while (C) and (D) correspond to a hydrophilic surface presence. Results for the wild-type (WT) variant are shown in green, and for Omicron in orange.

Now, we focus on the glycan-protein interaction. Contact maps for each of the monosaccharides in the N343 glycan and the protein residues in the contact region for WT (top) and omicron (bottom), plus the two model surfaces are shown in Fig. 8. They display the ratio between the number of frames along the trajectory where the contact is present and the total trajectory frames. All contact maps include a green dashed line separating the Head and RBM regions. The general difference between closed-head and closed-RBM conformations is that the glycan at the head region interacts mostly with the protein residues in this region (Figs. 8A,C, E, and G). This can be partly understood by the fact that the glycan attaches to the residue ASN 12 located at the head region.

**Figure 8.**
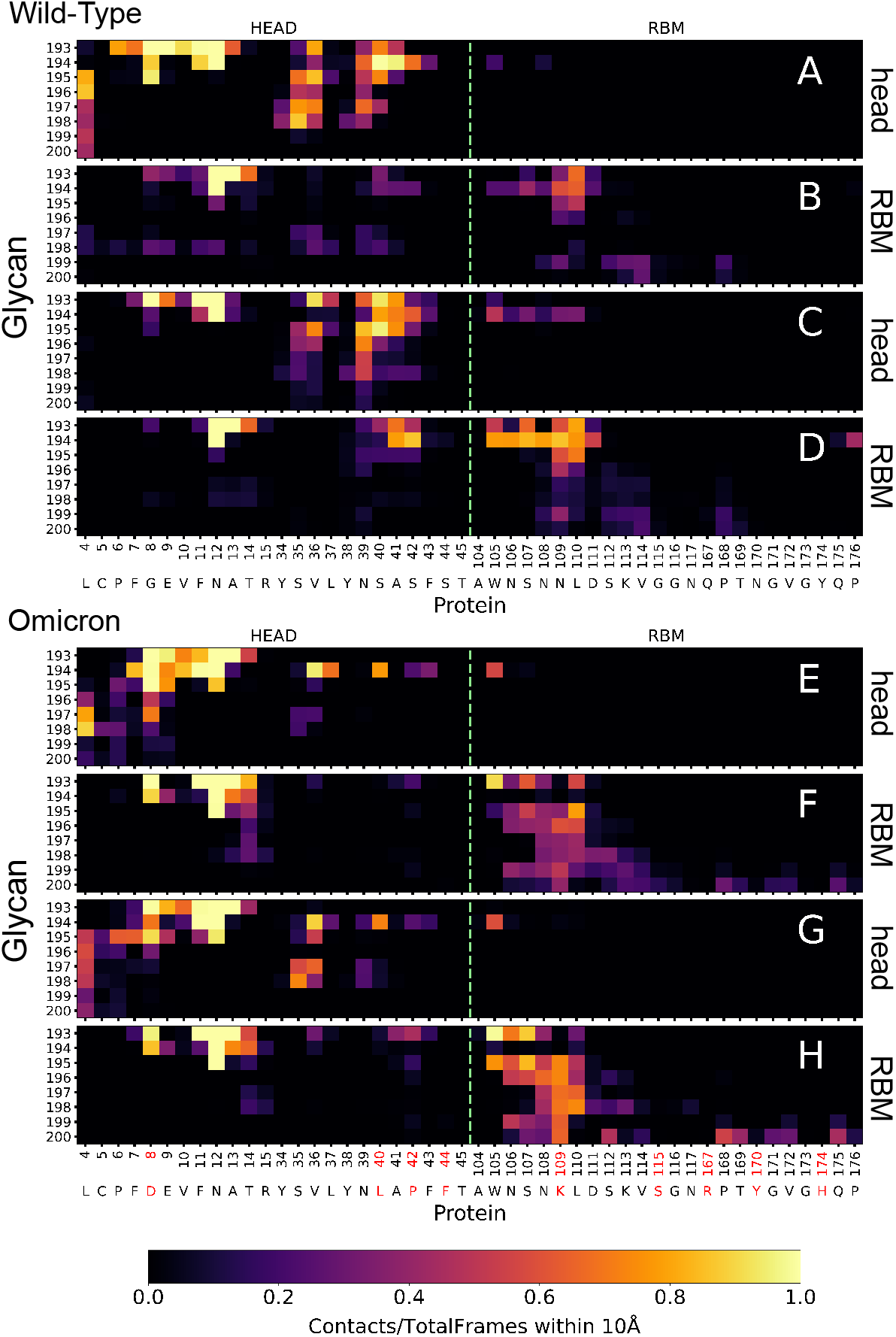
Contact maps between glycan and protein residues within the contact region for (A–D) WT and (E–H) Omicron. Panels (A, B, E, F) show results from simulations with a hydrophobic surface, while (C, D, G, H) correspond to a hydrophilic surface. In panels (A, C, E, G), the glycan is positioned between the RBD head and the surface; in (B, D, F, H), it is located between the receptor-binding motif (RBM) and the surface. A horizontal segmented lime green line in each map indicates the boundary separating residues in the RBD head from those in the RBM region of the RBD. Contacts are considered as residue– residue distances under 10 Å.

Hence, even for the closed-RBM conformation, the glycan has necessarily to interact with parts of the head region, mainly the protein fragment between residues 7-15. This can be clearly seen for omicron in Figs. 8F and H. Such tendency is not as strong for WT, which we attribute to the larger fluctuations of the glycan on the surface and towards the head region, as shown in pastel-red footprints (Fig. 4C and Fig. S2C). The closed-RBM conformations also show key differences between WT and Omicron due to the mutations (marked in red in the x-axis) that make omicron’s RBM more hydrophilic and lead to an increase in the total number of contacts and to extend them to all the protein residues in positions 104 to 115 (Figs. 8F and H). WT, without those additional hydrophilic residues, has a much narrower distribution of contacts (Figs. 8B and D). The later panel D shows a very strong interaction with AFUC 194 which we presume is an effect of the protein oscillatory adsorption on the hydrophilic surface.

Table 1 quantifies the number of contacts (distances below 10 Å) taking into account the electrostatic character of the residues (positively (CP) and negatively (CN) charged, non-polar (NP) and uncharged polar (UP)). Values are normalized to the product of the total number of simulation frames and the total number of glycan residues. Numbers in parentheses correspond to the total number of protein residues in each of the four categories. In the case of the glycan, we have three positively charged monosaccharides, two negatively charge and three neutral ones.

**Table 1:** Contact frequencies between protein residues in the contact region and the glycan, grouped by residue type (positively charged, negatively charged, non-polar, and uncharged polar) for WT and Omicron with a 10Å contact cutoff. Values are normalized by the total number of glycan residues, and the number of simulation frames considered. A value equal to the number of residue in a given type (in parenthesis) corresponds to the theoretical case where there is at least one contact of glycan residue with one protein residue. Note that glycan residue can have more than one contact with protein residues. Glycines have been excluded from the count due to their low polarity.

|  |  | Variant | Positive | Negative | Non-Polar | Uncharged Polar |
| --- | --- | --- | --- | --- | --- | --- |
| head | PBL0 | WT | 0.000(2) | 0.538(2) | 0.826 (18) | 0.624 (21) |
|  |  | Omicron | 0.002 (4) | 2.484 (3) | 0.666 (21) | 0.253 (16) |
|  | PBL1 | WT | 0.000 | 0.364 | 0.578 | 0.667 |
|  |  | Omicron | 0.008 | 1.487 | 0.710 | 0.363 |
| RBM | PBL0 | WT | 0.310 | 0.516 | 0.398 | 0.423 |
|  |  | Omicron | 0.937 | 1.053 | 0.455 | 0.817 |
|  | PBL1 | WT | 0.337 | 0.445 | 0.498 | 0.692 |
|  |  | Omicron | 1.237 | 0.876 | 0.552 | 0.962 |

Starting at the closed-head conformation on the hydrophobic surface (PBL0), in WT glycan contacts with negative residues are clearly favored: the average number of contacts with the two negative residues is of the same order as the ones with 21 uncharged-polar and 18 non-polar residues. For omicron, negative contacts completely dominate. This is clearly a collective effect that goes beyond the presence of one more negatively charged residue (ASP 8), and is probably due to the presence of two adjacent negatively charged residues (ASP 8 and GLU 9) located very closely to the glycan attachment location (ASN 12). The same trends are found on the hydrophilic surface.

The closed-RBM conformations show a dramatic increase in the number of contacts with positively charged (CP) residues. Again, this effect is more pronounced in Omicron and goes beyond the change in the total number of CP residues (2 in WT, 4 in Omicron), as previously discussed in other works focused on the entire RBD electrostatic interactions.^24,30,31^ The neighboring location of the LYS109 and LYS113 probably contributes to this enhancement, which is more pronounced in the hydrophilic surface. Moreover, the mutation to negatively charged ASP 8 in Omicron (former neutral GLY 8 in WT) shows strong interactions with the positively charged AFUC 194 that is very close to the anchoring monosaccharide (see the fluctuating areas of the glycans in Figures 5D and S7D). The conformation of the glycan at the RBM position could be also screening the contact with negative residues for Omicron. Contacts for neutral residues decrease for non-polar residues and significantly increase with uncharged polar ones in Omicron. These results suggest that the local charge distribution of the glycan is key to understand the dynamics and the interactions of glycans, and not only the globally neutral character of this polysaccharide. As it is the case for purely proteinaceous systems.^32^ Similar results are found in the analysis of contact maps where the cutoff distance has been extended to 15 Å(SI Fig. S11 and Table S8) and for the delta variant (SI Fig. S12 and Table S9).

### Upscaling of the RBD to the spike S1 glycoprotein onto polarizable surfaces

In order to validate our minimal model centered on the spike protein RBD, we extended our study to the complete S1 spike protein. Going beyond the small system has twofold objectives, first to verify that the initial conformations of the glycan at the RBD we assumed (closed-head and closed-RBM) are found in company of the up, down, down (UDD) spike conformation. In Fig. 9, we analyze the 2D-densities as contours of the bottom RBDs trimer comprising the spike protein and quantify the number of contacts made by the S1 spike at the contact regions studied in our small RBD centered system. For the hydrophobic surface shown in Figs. 9A and B, we recognize that the contour areas of the small system (dashed lines) are mostly contained within the S1 spike system (solid lines). The number of contacts are coinciding with the residues focused in our studies by 80*±*10%. The behavior of the glycoprotein onto the hydrophilic surface (Figs. 9C and D) shows also a coincidence of contour areas but in lesser amount than the hydrophobic surface. Similarly, with the contacts, which match mostly the residues between the range of 100-175. Regarding the coincidence of glycan conformations, we quantified the number of contacts between the RBD Glycans and the proteins from the whole S1 spike simulations in Table 2. Strikingly, the conformations we studied at the RBD level for the closed RBD add-up more than half of total interactions between the protein and glycans at the S1 level for WT and Omicron and in both conformations open and closed (see Table 2).

**Figure 9.**
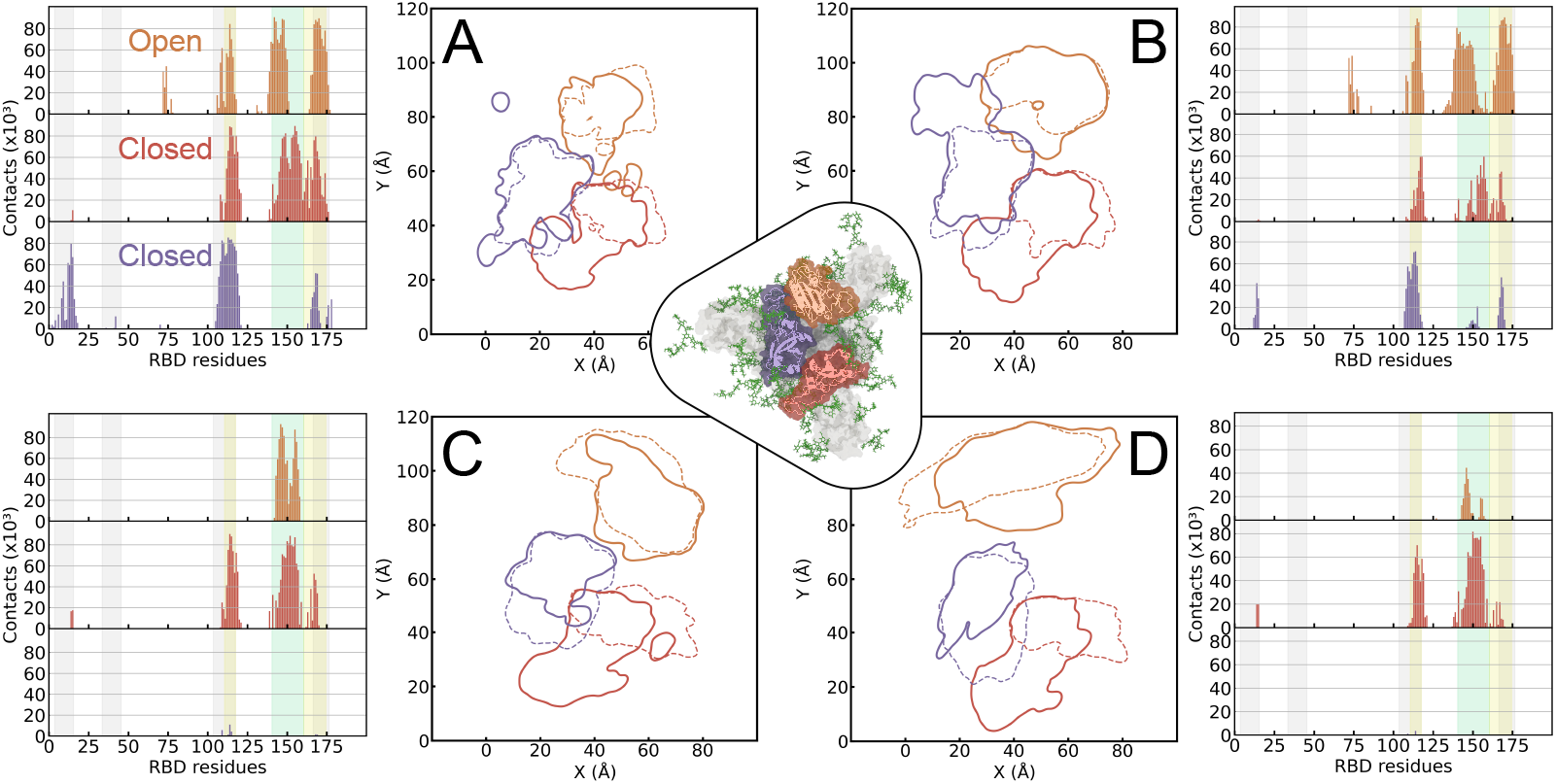
RBDs in an open–closed–closed S1 Spike protein configuration. Each panel shows a contour plot representing the contact regions of the three Spike RBDs in the presence of PBLs, and contact histograms of the RBD residues. In the contour plots, dashed lines indicate the contact regions analyzed in the isolated-RBD simulations, and solid lines enclose the top 50 residues with the most surface contacts. In the contact histograms, green and orange regions highlight the two contact regions previously considered for the open-RBD configuration. ^8^ In contrast, gray highlights the contact region of the closed configuration analyzed in this manuscript. RBD colors are consistent across the plots, with the dark orange RBD corresponding to the open configuration. Panels (A) and (B) show the results for the WT variant in the presence of hydrophobic and hydrophilic surfaces, respectively. Panels (C) and (D) correspond to the Omicron variant under the same conditions. In the center of the figure, there is a representation of the S1 glycoprotein.

**Table 2:**
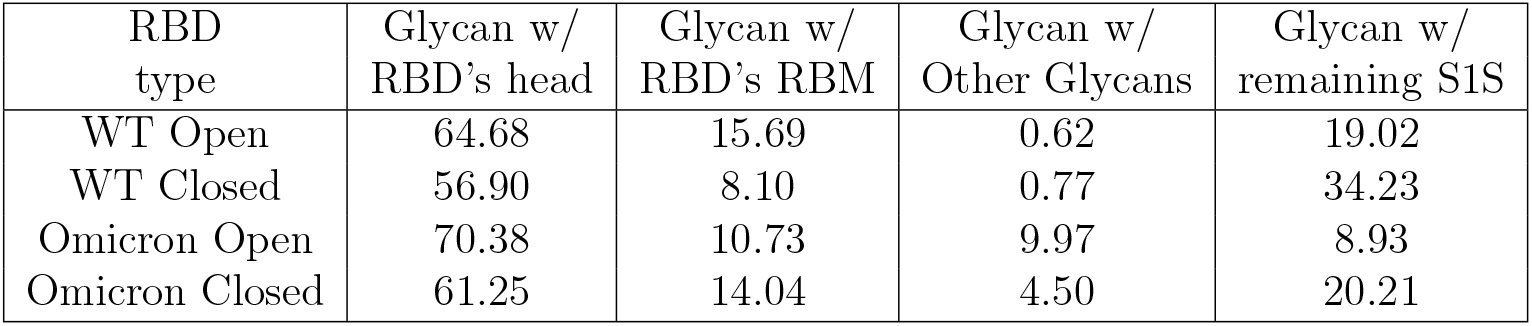
Glycan interactions for S1 glycoprotein–PBLs simulations. Table reports the percentual contribution of the contacts with the head and RBM region of RBD it is attached to, with other glycans and the remaining protein residues of the S1 glycoprotein. Note that the snapshots of each system are found in the SI Figures S13, S14, S15 and S16

| RBD type | Glycan w/ RBD's head | Glycan w/ RBD's RBM | Glycan w/ Other Glycans | Glycan w/ remaining S1S |
| --- | --- | --- | --- | --- |
| WT Open | 64.68 | 15.69 | 0.62 | 19.02 |
| WT Closed | 56.90 | 8.10 | 0.77 | 34.23 |
| Omicron Open | 70.38 | 10.73 | 9.97 | 8.93 |
| Omicron Closed | 61.25 | 14.04 | 4.50 | 20.21 |

The rest of interactions of the N343 glycan have been tracked with other regions of the glycoprotein complex (besides their corresponding RBD contact regions either head or RBM), namely, the non-RBD S1 protein residues, the non-RBD other Glycans (beyond the N343 one placed at the RBD site). Remarkably those 2 interactions do not exceed a 20% of contacts for the S1 UDD conformation. Another interesting result is that the closed-head conformation is at least 2 times more frequent than the closed-RBM all at the S1 spike level. This is partly tracked back to the fact that the Glycan is attached in the head region. However, it is also related to the availability of the contact regions, as we saw that the proteinaceous contact frequencies for the S1 complex are mainly located at the RBM site. In other words, it turns a busy flag onto this site, while the head region contains mostly fewer contacts (see Fig. 9. Another interaction that has been made tractable within this analysis is the interaction of other glycans of the spike S1 (WT or Omicron, see SI Table S10) with either the head and/or RBM regions which represent a maximum of ≈ 20% for WT and ≈ 30% for omicron. The external glycans also have a clear preference to interact with the head site of both variants and any conformation. Some snapshots illustrating the interactions of the ‘glycan context’ are shown in the SI Figs. S13, S14, S15 and S16.

## Discussion

We studied the role of glycans in the adsorption of RBD glycoproteins onto planar hydrophobic and hydrophilic surfaces. We use model surfaces in order to generalize our results to specific short-range interactions, which captures the effect of mutations on different RBD glycoprotein VoCs at their contact regions. Our investigation highlights the glycans’ dual capacity, to act either as a glue that dominates the whole RBD glycoprotein adsorption strength via different tethering patterns, or as a blocking element preventing the protein-surface interactions. The existence of several mutations in the RBD glycoproteins provides the perfect playground to spatially understand the effect of mutations in the contact regions via short-range interactions, particularly between the WT and Omicron variants. Here, a challenge is to identify the location of those interactions, which we attend with the help of our planar surface model, enabling the evaluation of those interactions with fewer degrees of freedom than a bulk representation. For the “on surface” analysis, we chose a combination of polymer adsorption theory^19–21^ with recent advanced 2D analysis,^23^ on top of specifically developed routines to evaluate the context of glycans and proteins during adsorption. These analyses allow us to identify and quantify the rich phenomenology of a highly flexible molecule upon adsorption. The deformation (ratio 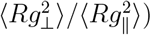 of the glycoproteins on the surface shows that the glycan can substitute the flexibility of the region exposed to the surface of a proteinaceous fragment at the glycoprotein-surface interface. We showed, in previous work for the open RBDs,^8^ that flexibility at such an interface can promote the number of contacts and increase the deformability of the glycoprotein upon adsorption. Whereby the role of the glycan for the open RBDs adsorption was minor, however, in the closed conformation, the glycan shows a dominant character upon adsorption. This is shown by the capacity to enable or hinder the effects of flexibility in different protein contact regions and, hence, influence the whole interaction at the RBD-surface interface. In particular, we show that the RBM contact-region is up to 50% more flattened (Fig. S1) upon adsorption than the head one, and thus blocking those specific short-range interactions at the RBM region clearly reduces the number of contacts at its interface. Interestingly, mutations are another parameter that also plays a key role in the deformation during adsorption, because depending on the character of the residues, they will improve selectively and/or collectively hydrophobic and hydrophilic interactions with each of the tested surfaces. We expect that the selective initial conformation of glycans towards the protein wall could be experimentally controlled as shown by adsorption studies onto different surfaces, ^33^ in this case however, the local electrostatic properties of each monosaccharide could be exploited for the functionalization of glycans on top of protein mutations.

Our results based on the initial position of the glycan at either closed-head or closed-RBM regions also provided very detailed observations, which we discuss in the following paragraphs along with the evaluated interfacial properties. The contact area in the closed-head conformation at least doubles that of the closed-RBM for the hydrophilic surfaces and all VoCs. Another quantity we computed is the footprints of the top residues with most contacts, showing the residues with stronger interactions of the protein residues, and at the same time allow us to interpret the glycan preferences in the ‘sandwich’ context between the RBD protein and the planar hydrophobic or hydrophilic surface. This analysis helped understand three archetypal behaviors of the glycan upon adsorption, namely: tethered adsorption, repulsive squeezing, and the soft tethering (intermittent anchoring). These captured behaviors can be combined dynamically during adsorption depending on the precise glycan environment. For instance, driven by mutations in the omicron head favoring hydrophobic interactions, the protein is able to squeeze-out the glycan over the hydrophobic surface, where the glycan faces a new interaction context at the liquid-surface interface. Similarly, the WT RBD glycan in the closed-RBM conformation is able to tether the hydrophobic surface and find strong hydrophobic interactions that can hold the rest of the glycoprotein during desorption processes. Similar observations are currently discussed in the literature for glycan-membrane^34,35^ interactions, for example, and glycan-glycan and glycan-protein^13,36^ ones. However to the best of our knowledge, our work is pioneering in the field of characterizing glycan properties in ‘2D’ during adsorption processes. Another analyzed quantity shows the footprints of the residues and glycan with top RMSFs, which allows to identify whether fluctuations promote more contacts at the interface or those contacts are rather based on highly localized short-range interactions with less flexible protein regions. For example, WT in the closed-RBM conformation has 3 common residues between the highest contacts and fluctuations at the interface. Moreover, the same glycan shows high flexibility and large fluctuations reaching two orders of magnitude greater values than those of the highest-fluctuating protein residues, allowing a “squeeze out” and tethering phenomena, as previously described. The amphiphatic character of the RBD’s glycan is also analyzed conveniently with our model surface setup. Here, we can proof that for this specific and globally neutral glycan, its hydrophobic number of contacts is constantly higher than the hydrophilic ones towards each corresponding surface. This was also confirmed by computing the angles of the sugar rings with respect to the surface normal.

A challenge we also tackled was the upscaling of our single-RBDs results to the whole trimer context (S1 fragment of the spike glycoprotein), given that the spike protein structural models have been well studied and shared within the computational biophysical community.^37,38^ The systematic upscaling study successfully validates the extended simulations of the RBD-only system by strikingly matching over half of the interactions with our closed-head and closed-RBM initial conformations.

In a nutshell, we have presented different landing footprints and deformation mechanisms of the RBD glycoproteins, which can be directly applied to understand the influence of the glycans on their adsorption. Beyond providing molecular understanding of the complex interplay between molecules of different flexibilities, namely, protein fragments and glycans. This work offers a systematic method to analyze glycoproteins short-range interactions, which at the same time can provide proper molecular interpretation of adsorption experiments with high-resolution experimental techniques.^11,25,29^

The novel adsorption mechanisms identified between glycoprotein and convenient model surfaces together with the very recent development of methods that allow tracking glycans with angstrom resolution,^29^ among others, provide the starting point and momentum to molecular understand these processes at heterogeneous interfaces and systematically look at applied systems, such as protein aggregation, multi-protein assemblies, misrecognition of protein or surfaces functionalized with glycans.

## Methods

### Molecular Dynamics

For this study, we have computed simulations of different scales of the system, the receptor binding domains, and S1 Spike protein of SARS-CoV-2. In particular, we have designed a model for the RBD that mimics the closed-RBDs in the trimeric S1 glycoprotein, adding its corresponding Glycan according to Casalino et. al, ^1^ and restraining the residues in the S1-RBD interface to mimic the presence of the rest of the Spike protein. On the other hand, configurations of the WT and Omicron open-S1 glycoprotein were also adopted from Casalino et. al.^1^ All-atom simulations were carried out with Gromacs 2023^39^ and the system components (protein, ions, and the polarizable bilayer) were modeled using CHARMM36^40,41^ forcefield, and TIP3P^42^ for the water. CHARMM-GUI was used to join the glycan to the RBDs. Energy minimization used CPUs, while all production runs used 1xGPU, as the former scaled better than 2xGPUs for our systems. The hydrophobic (PBL0) and hydrophilic (PBL1) surfaces were built from a small patch of decanol (DOL), in which restraints were used in order to maintain the bilayer shape and avoid any effects of the mechanical properties of the surface, as discussed in previous works,^22,43,44^ the bilayer model does not include any curvature(is not flexible) and also no molecular defects. The replicated PBL (using *gmx editconf*) patch was solvated in a slab-formed water box. The RBD models for WT, Delta, and Omicron were added to cubic boxes (see 3) containing the polarized bilayers. The OH-groups of the DOL chains were tuned to 0 or 1 for hydrophobic and hydrophilic surfaces, respectively. ^22^ This procedure was repeated 3 times for each polarity, aggregating from 3 replicas per VoCs (WT, delta, and omicron), and 2 glycan configurations relative to the RBD, obtaining a total of 18 configurations. All simulations included ions to work under neutral charge conditions. Periodic boundary conditions were applied, and PME was used for long-range electrostatics. Minimization was done by steepest descent (50000 steps) with integration steps of 0.01 ps. The equilibration time for the NVT and NPT was 100ps, respectively. For each isolated RBD simulation, we determined the contacts between the RBD and the S1 region. Those contacts were applied as position restraints of 250 kJ mol^-1^ nm^-2^ in x and y axes. In other words, we considered as flexible regions of the RBD all the others that are not in contact with the S1. However, in order to quantify adsorption, we kept the z-axis free of restraints in all RBD-S1 contacts. Note also that the position restraints were not added to residues in the contact region. Mechanical restraints were not added to simulations of the entire S1 glycoprotein, as it is bound to the highly flexible S2 membrane protein, which is simultaneously shown to be highly diffusive over the envelope.^10,45,46^ Production simulations began from the final equilibrated snapshots, and three copies (with angular rotations of up to 3 degrees from the reference) of each system were simulated. Finally, 300 ns of trajectories of each replica were collected as described in Table 3.

**Table 3.** The table contains the general configurations of the simulated systems. Note that 848 decanol molecules were used in simulations of the isolated RBDs in the presence and absence of glycans, and 5028 decanol molecules in simulations of the whole S1 Spike protein. Simulations of RBD-PBLs with glycans are performed in two different configurations of the glycans relative to the RBD: a glycan in the head-surface interface (head) and RBM-surface interface (RBM) configuration. Representations of the initial configurations are shown in Fig.1.

|  | VoC | No. of replicas | Box size | Atoms | Time per replica |
| --- | --- | --- | --- | --- | --- |
| RBD | WT | 3 | $7.5 \times 7.5 \times 12$ | 2980 | 300 |
| | Delta | 3 | $7.5 \times 7.5 \times 12$ | 2993 | 300 |
| | Omicron | 3 | $7.5 \times 7.5 \times 12$ | 3036 | 300 |
| | WT+Glyc | 6 (3 head + 3 RBM) | $8.6 \times 7.6 \times 12$ | 2979+192 | 300 |
| | Delta+Glyc | 6 (3 head + 3 RBM) | $8.6 \times 7.6 \times 12$ | 2992+192 | 300 |
| | Omicron+Glyc | 6 (3 head + 3 RBM) | $8.6 \times 7.6 \times 12$ | 3035+192 | 300 |
| Spike | WT | 3 | $16.9 \times 22.5 \times 25.0$ | 63793 | 300 |
| | Omicron | 3 | $16.9 \times 22.5 \times 25.0$ | 64339 | 300 |

Note that all production runs used an integration step of 2 fs. Starting configurations from the MD production can be found in a Zenodo repository Our analyses were carried out using our previously published MDAnalysis-based toolkit 2DAnalysis, ^23^ core MDAnalysis functions, and in-GROMACS tools such as SASA computation.

### Density contour plots

Density contour plots were generated with the 2DAnalysis toolkit^23^ and applied to footprint residue figures (Figures 4, 5, S2, S3,S7, S8) and hydrogen-bond figures (Figures 6, S9) to map the positioning of the protein, and the glycan during simulations. Only residues within contact regions and frames where the contact regions of the protein were within 15 Å of decanol oxygen atoms were included, excluding desorbed frames.

For Figures 5, S7 and S8, residue areas were obtained by integrating KDE contour densities of each residue during the simulation using the getArea function in 2DAnalysis. In these cases, the contour densities of residues were computed considering the 1500 (300) frames with the most contact of the protein’s contact region to hydrophobic (hydrophilic) surface to ensure comparable areas for different replicas, variants, and glycan positioning.

### Protein-Glycan Contact Maps

Contact Map was computed by measuring the distance between the centers of mass of each residue in the contact region of the protein and glycan. All distances under 10Å between the residues of these two groups were considered a contact. Figures S11, and S12, show contact maps considering distances under 15Å.

## Author Contributions

H.V.G. conceived the research; H.V.G., A.M.B.-F and R. P. designed the research; A.M.B.- F performed the molecular dynamics simulations; H.V.G., A.M.B.-F, and W.M. performed basic and advanced analysis; H.V.G., A.M.B.-F and R. P. interpreted the data; R.P. and H.V.G. wrote the manuscript and supervised the research. All authors have read and agreed to the published version of the manuscript.

## Acknowledgement

We thank Matej Kanduč for illuminating discussions on the molecular modeling of polarizable surface, Dr. Abigail Dommer for advice regarding the modeling of the whole spike trimer with glycans. H.V.G. acknowledges financial support from the Ramón y Cajal grant No. RYC2022-038082-I and Spanish Ministry of Science and Innovation, through project PID2023-150536NA-I00, and the “Severo Ochoa” Grant No. CEX2023-001263-S for Centers of Excellence; and CSIC’s grant MMT24-ICMAB-01 for the nanoML4Med project. H.V.G. acknowledges also Red Española de Supercomputación (RES) for the computing time and technical support at the Finisterrae III supercomputer projects FI-2025-2-0062 and FI-2025- 2-0058. R.P. acknowledges support from the Spanish Ministry of Science and Innovation, through project PID2020-115864RB-I00 and the “María de Maeztu” Programme for Units of Excellence in R&D (CEX2018-000805-M). A.M.B.-F. acknowledges support from the autonomous community of Madrid, through the contract PIPF-2023/TEC-29920.

## Supplementary information

The Supporting Information available within this article contains: Tables indicating the groups in each group, 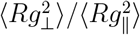 ratios and their corresponding ⟨*Rg*_⊥_⟩ and ⟨*Rg*_∥_⟩ contributions, and % H-Bond during simulations; and figures show the 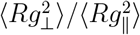 per region, Contour plots with footprint of top 5 Contacts and Top 5 RMSFs for Delta, and for WT and Omicron onto hydrophilic surfaces, representative snapshots of RBD interactions to the surfaces, violin plot of the relative angles of the sugar rings of the glycans with the normal of the surface, % H-Bond during simulations for Delta, the Contact Maps for Delta with a 10Å threshold, and WT, delta and Omicron with 15Å, and snapshot of the context of glycan within the S1 Spike glycoprotein.

## Data and Software Availability

All-atom simulations were carried out with Gromacs 2023; the corresponding parameter files, input files, topologies, position restraints, and initial configurations are available on the Zenodo repository (https://zenodo.org/records/17207608), and analysis scripts can be found in the Github repositories https://github.com/pyF4all/2DanalysisTutorials and https://github.com/ABoschF/Paper_Closed-RBDs_public.

