## Supplementary material for "Glycans modulate the adsorption of RBD Glycoproteins on polarizable surfaces": SI

‖*Biophysics and Intelligent Matter Lab, E-08193 Barcelona, Spain*

### Supplementary Tables

Table S1: $\langle Rg^2_{\perp}\rangle/\langle Rg^2_{\parallel}\rangle$ radii of gyration ratios of the contact region of the closed and open RBDs, and the glycan during the simulations of the closed RBDs with the two glycan-RBD conformations for the three VoCs. These values are represented as symbols in Figure 2. Ratio values under 0.32 are considered to be in the polymer semiflexible regime.[1] Open does not have closed/head or closed-RBM configurations; its contact region is only proteic.[2]

| | Glycan | Variant | $\langle Rg^2_{closed,\perp}\rangle/\langle Rg^2_{closed,\parallel}\rangle$ | $\langle Rg^2_{glycan,\perp}\rangle/\langle Rg^2_{glycan,\parallel}\rangle$ | $\langle Rg^2_{open,\perp}\rangle/\langle Rg^2_{open,\parallel}\rangle$ |
|---|---|---|---|---|---|
| PBL0 | head | WT | 0.071 | 0.087 | 0.060 |
| | | Delta | 0.081 | 0.119 | 0.057 |
| | | Omicron | 0.065 | 0.130 | 0.064 |
| | RBM | WT | 0.105 | 0.149 | - |
| | | Delta | 0.080 | 0.060 | - |
| | | Omicron | 0.072 | 0.183 | - |
| PBL1 | head | WT | 0.070 | 0.385 | 0.060 |
| | | Delta | 0.074 | 0.258 | 0.057 |
| | | Omicron | 0.077 | 0.205 | 0.060 |
| | RBM | WT | 0.103 | 0.445 | - |
| | | Delta | 0.079 | 0.368 | - |
| | | Omicron | 0.070 | 0.318 | - |

Table S2: Mean perpendicular and parallel radii of gyration of the contact region of the closed and open RBDs, and the glycan during the simulations of the closed RBDs with the two glycan-RBD conformations for the three VoCs. These values are represented as symbols in Figure 2. Open does not have closed/head or closed-RBM configurations; its contact region is only proteic.[2]

| | | Variant | $Rg_{\perp}^{closed}$ | $Rg_{\parallel}^{closed}$ | $Rg_{\perp}^{glyc}$ | $Rg_{\parallel}^{glyc}$ | $Rg_{\perp}^{open}$ | $Rg_{\parallel}^{open}$ |
|---|---|---|---|---|---|---|---|---|
| PBL0 | head | WT | 3.43 | 12.87 | 2.23 | 7.71 | 3.64 | 14.85 |
| | | Delta | 3.83 | 13.46 | 2.50 | 7.37 | 3.59 | 15.04 |
| | | Omicron | 3.44 | 13.47 | 2.51 | 7.45 | 3.84 | 15.14 |
| | RBM | - WT | 4.12 | 12.72 | 2.64 | 7.09 | - | - |
| | | Delta | 3.80 | 13.48 | 1.87 | 7.85 | - | - |
| | | Omicron | 3.65 | 13.58 | 2.82 | 7.10 | - | - |
| PBL1 | head | WT | 3.43 | 12.95 | 4.02 | 6.66 | 3.59 | 14.70 |
| | | Delta | 3.66 | 13.51 | 3.34 | 6.91 | 3.58 | 15.04 |
| | | Omicron | 3.63 | 13.10 | 3.01 | 7.07 | 3.74 | 15.30 |
| | RBM | WT | 4.08 | 12.73 | 4.10 | 6.33 | - | - |
| | | Delta | 3.74 | 13.37 | 3.95 | 6.66 | - | - |
| | | Omicron | 3.55 | 13.39 | 3.59 | 6.78 | - | - |

Table S3: Table contains the ResIDs and corresponding residue names of residues used in Group 1 (Head of the RBD). Mutations are highlighted in fuchsia. **CN**: Charged Negative, **CP**: Charged Positive, **UP**: Uncharged Polar and **NP**: NonPolar

| | ResIDs | WT | Delta | Omicron |
|---|---|---|---|---|
| 1 | 4 | LEU (NP) | LEU (NP) | LEU (NP) |
| 2 | 5 | CYS (NP) | CYS (NP) | CYS (NP) |
| 3 | 6 | PRO (NP) | PRO (NP) | PRO (NP) |
| 4 | 7 | PHE (NP) | PHE (NP) | PHE (NP) |
| 5 | 8 | GLY (NP) | GLY (NP) | **ASP (CN)** |
| 6 | 9 | GLU (CN) | GLU (CN) | GLU (CN) |
| 7 | 10 | VAL (NP) | VAL (NP) | VAL (NP) |
| 8 | 11 | PHE (NP) | PHE (NP) | PHE (NP) |
| 9 | 12 | ASN (UP) | ASN (UP) | ASN (UP) |
| 10 | 13 | ALA (NP) | ALA (NP) | ALA (NP) |
| 11 | 14 | THR (UP) | THR (UP) | THR (UP) |
| 12 | 15 | ARG (CP) | ARG (CP) | ARG (CP) |
| 13 | 34 | TYR (UP) | TYR (UP) | TYR (UP) |
| 14 | 35 | SER (UP) | SER (UP) | SER (UP) |
| 15 | 36 | VAL (NP) | VAL (NP) | VAL (NP) |
| 16 | 37 | LEU (NP) | LEU (NP) | LEU (NP) |
| 17 | 38 | TYR (UP) | TYR (UP) | TYR (UP) |
| 18 | 39 | ASN (UP) | ASN (UP) | ASN (UP) |
| 19 | 40 | SER (UP) | SER (UP) | **LEU (NP)** |
| 20 | 41 | ALA (NP) | ALA (NP) | ALA (NP) |
| 21 | 42 | SER (UP) | SER (UP) | **PRO (NP)** |
| 22 | 43 | PHE (NP) | PHE (NP) | PHE (NP) |
| 23 | 44 | SER (UP) | SER (UP) | **PHE (NP)** |
| 24 | 45 | THR (UP) | THR (UP) | THR (UP) |

Table S4: Table contains the ResIDs and corresponding residue names of residues used in Group 2 (Contact-to-surface RBM region). Mutations are highlighted in fuchsia. **CN**: Charged Negative, **CP**: Charged Positive, **UP**: Uncharged Polar and **NP**: NonPolar

| | ResIDs | WT | Delta | Omicron |
|---|---|---|---|---|
| 1 | 104 | ALA (NP) | ALA (NP) | ALA (NP) |
| 2 | 105 | TRP (NP) | TRP (NP) | TRP (NP) |
| 3 | 106 | ASN (UP) | ASN (UP) | ASN (UP) |
| 4 | 107 | SER (UP) | SER (UP) | SER (UP) |
| 5 | 108 | ASN (UP) | ASN (UP) | ASN (UP) |
| 6 | 109 | ASN (UP) | ASN (UP) | **LYS (CP)** |
| 7 | 110 | LEU (NP) | LEU (NP) | LEU (NP) |
| 8 | 111 | ASP (CN) | ASP (CN) | ASP (CN) |
| 9 | 112 | SER (UP) | SER (UP) | SER (UP) |
| 10 | 113 | LYS (CP) | LYS (CP) | LYS (CP) |
| 11 | 114 | VAL (NP) | VAL (NP) | VAL (NP) |
| 12 | 115 | GLY (NP) | GLY (NP) | **SER (UP)** |
| 13 | 116 | GLY (NP) | GLY (NP) | GLY (NP) |
| 14 | 117 | ASN (UP) | ASN (UP) | ASN (UP) |
| 15 | 167 | GLN (UP) | GLN (UP) | **ARG (CP)** |
| 16 | 168 | PRO (NP) | PRO (NP) | PRO (NP) |
| 17 | 169 | THR (UP) | THR (UP) | THR (UP) |
| 18 | 170 | ASN (UP) | ASN (UP) | **TYR (UP)** |
| 19 | 171 | GLY (NP) | GLY (NP) | GLY (NP) |
| 20 | 172 | VAL (NP) | VAL (NP) | VAL (NP) |
| 21 | 173 | GLY (NP) | GLY (NP) | GLY (NP) |
| 22 | 174 | TYR (UP) | TYR (UP) | **HIS (CP)** |
| 23 | 175 | GLN (UP) | GLN (UP) | GLN (UP) |
| 24 | 176 | PRO (NP) | PRO (NP) | PRO (NP) |

Table S5: Residues involved in hydrogen bonding during simulations of the Wild-Type (WT) variant in three configurations: open RBD, closed with the glycan positioned between the RBD head and the surface, and closed with the glycan between the RBM and the surface (top to bottom, respectively). The table reports the residue IDs, names, and the percentage of simulation time in which each residue participated in hydrogen bonding. Top five residues are represented in Fig. 6.

| | ResIDs | ResNames | % Hbonds |
|---|---|---|---|
| WT Glycan in Head | 197 | AMAN | 36.24 |
| | 199 | AMAN | 35.80 |
| | 198 | BGLCNA | 30.64 |
| | 200 | BGLCNA | 22.40 |
| | 194 | AFUC | 9.64 |
| | 196 | BMAN | 6.44 |
| | 195 | BGLCNA | 4.56 |
| | 109 | ASN | 3.16 |
| | 39 | ASN | 0.13 |
| | 38 | TYR | 0.02 |
| WT Glycan in RBM | 199 | AMAN | 17.42 |
| | 200 | BGLCNA | 14.84 |
| | 197 | AMAN | 14.07 |
| | 198 | BGLCNA | 10.04 |
| | 39 | ASN | 7.53 |
| | 195 | BGLCNA | 5.64 |
| | 196 | BMAN | 2.49 |
| | 38 | TYR | 2.31 |
| | 35 | SER | 0.91 |
| | 194 | AFUC | 0.78 |
| | 42 | SER | 0.04 |
| WT OPEN | 169 | THR | 54.67 |
| | 167 | GLN | 17.90 |
| | 153 | GLU | 11.20 |
| | 115 | GLY | 1.63 |
| | 118 | TYR | 0.83 |
| | 170 | ASN | 0.60 |
| | 116 | GLY | 0.57 |
| | 147 | THR | 0.43 |
| | 117 | ASN | 0.33 |
| | 150 | ASN | 0.27 |
| | 156 | ASN | 0.07 |
| | 146 | SER | 0.03 |

Table S6: Residues involved in hydrogen bonding during simulations of the Omicron variant in three configurations: open RBD, closed with the glycan positioned between the RBD head and the surface, and closed with the glycan between the RBM and the surface (top to bottom, respectively). The table reports the residue IDs, names, and the percentage of simulation time in which each residue participated in hydrogen bonding. Top five residues are represented in Figure 6.

| | ResIDs | ResNames | % Hbonds |
|---|---|---|---|
| Omicron Glycan in Head | 194 | AFUC | 54.31 |
| | 199 | AMAN | 48.40 |
| | 109 | LYS | 35.00 |
| | 197 | AMAN | 29.98 |
| | 200 | BGLCNA | 27.16 |
| | 39 | ASN | 20.58 |
| | 198 | BGLCNA | 20.58 |
| | 196 | BMAN | 14.27 |
| | 195 | BGLCNA | 8.67 |
| | 193 | BGLCNA | 7.89 |
| | 108 | ASN | 1.33 |
| | 169 | THR | 0.16 |
| | 38 | TYR | 0.04 |
| Omicron Glycan in RBM | 200 | BGLCNA | 60.62 |
| | 199 | AMAN | 44.44 |
| | 197 | AMAN | 41.71 |
| | 196 | BMAN | 39.53 |
| | 39 | ASN | 17.73 |
| | 194 | AFUC | 15.53 |
| | 195 | BGLCNA | 10.67 |
| | 198 | BGLCNA | 8.09 |
| | 38 | TYR | 0.87 |
| | 109 | LYS | 0.16 |
| Omicron Open RBD | 170 | TYR | 48.97 |
| | 167 | ARG | 32.73 |
| | 169 | THR | 14.53 |
| | 115 | SER | 13.80 |
| | 150 | ASN | 7.97 |
| | 156 | ASN | 5.60 |
| | 118 | TYR | 5.40 |
| | 147 | LYS | 0.70 |
| | 158 | TYR | 0.30 |
| | 154 | GLY | 0.23 |
| | 151 | GLY | 0.17 |

Table S7: Residues involved in hydrogen bonding during simulations of the Delta variant in three configurations: open RBD, closed with the glycan positioned between the RBD head and the surface, and closed with the glycan between the RBM and the surface (top to bottom, respectively). The table reports the residue IDs, names, and the percentage of simulation time in which each residue participated in hydrogen bonding. Top five residues are represented in Figure S9.

| | ResIDs | ResNames | % Hbonds |
|---|---|---|---|
| Delta Glycan in Head | 197 | AMAN | 42.02 |
| | 199 | AMAN | 35.47 |
| | 194 | AFUC | 32.82 |
| | 198 | BGLCNA | 31.62 |
| | 200 | BGLCNA | 29.07 |
| | 196 | BMAN | 11.33 |
| | 109 | ASN | 9.93 |
| | 195 | BGLCNA | 0.49 |
| | 113 | LYS | 0.42 |
| | 39 | ASN | 0.09 |
| | 108 | ASN | 0.07 |
| | 169 | THR | 0.02 |
| Delta Glycan in RBM | 200 | BGLCNA | 43.40 |
| | 197 | AMAN | 29.60 |
| | 199 | AMAN | 28.87 |
| | 198 | BGLCNA | 19.27 |
| | 196 | BMAN | 3.96 |
| | 194 | AFUC | 2.00 |
| | 195 | BGLCNA | 1.51 |
| | 39 | ASN | 0.24 |
| | 109 | ASN | 0.20 |
| | 113 | LYS | 0.02 |
| Delta OPEN | 153 | GLU | 46.80 |
| | 167 | GLN | 15.63 |
| | 115 | GLY | 10.93 |
| | 169 | THR | 4.47 |
| | 147 | LYS | 3.70 |
| | 118 | TYR | 3.27 |
| | 150 | ASN | 0.90 |

Table S8: Contact frequencies between protein residues and the glycan, grouped by residue type (positively charged, negatively charged, non-polar, and uncharged polar) for WT and Omicron with a 15Å contact cutoff. Values are normalized by the total number of glycan residues, and the number of simulation frames considered. A value equal to the number of residue in a given type (in parenthesis) corresponds to the theoretical case where there is at least one contact of glycan residue with one protein residue. Note that glycan residue can have more than one contact with protein residues. Glycines have been excluded from the count due to their low polarity.

| | | Variant | Positive | Negative | Non-Polar | Uncharged Polar |
|---|---|---|---|---|---|---|
| head | PBL0 | WT | 0.041 (2) | 1.712 (2) | 2.963 (18) | 1.805 (21) |
| | | Omicron | 0.265 (4) | 4.484 (3) | 2.430 (21) | 1.193 (16) |
| | PBL1 | WT | 0.191 | 1.701 | 2.362 | 2.126 |
| | | Omicron | 0.258 | 3.688 | 2.720 | 1.489 |
| RBM | PBL0 | WT | 2.584 | 2.981 | 1.756 | 1.650 |
| | | Omicron | 3.503 | 3.463 | 2.226 | 2.286 |
| | PBL1 | WT | 2.794 | 2.703 | 1.894 | 2.348 |
| | | Omicron | 3.505 | 3.561 | 2.682 | 2.729 |

Table S9: Contact frequencies between protein residues and the glycan, grouped by residue type (positively charged, negatively charged, non-polar, and uncharged polar) for Delta with 10Å and(|) 15Å cutoff, analogue treatment to tables 1 and S8. In parentheses, the number of residues of the corresponding residue type, and glycines have been excluded from the count due to their low polarity.

| | Surface | Variant | Positive | Negative | Non-Polar | Uncharged Polar |
|---|---|---|---|---|---|---|
| head | PBL0 | Delta | 0.000 \|0.009 (2) | 0.429 \|1.769 (2) | 0.915 \|3.017 (18) | 0.463 \|1.759 (21) |
| | PBL1 | Delta | 0.000 \|0.114 | 0.264 \|1.303 | 0.651 \|2.525 | 0.596 \|1.963 |
| RBM | PBL0 | Delta | 1.740 \|4.696 | 1.366 \|3.920 | 0.547 \|1.799 | 0.607 \|2.062 |
| | PBL1 | Delta | 0.316 \|1.821 | 0.538 \|2.419 | 0.570 \|1.936 | 0.555 \|1.962 |

Table S10: Glycan interactions for S1 glycoprotein–PBLs simulations. Table reports the mean contact occurrence of glycans during the simulations. ”Glycan” label refers to the Glycan in the corresponding RBD, ”Other Glycans” refers to all glycans but the glycan in the corresponding RBD, and ”remaining S1S” refers to all the protein residues in the S1 Spike protein, but those in the RBD to which it is attached.

| RBD type | Glycan w/ RBD's head | Other Glycans w/ RBD's head | Glycan w/ Other Glycans |
|---|---|---|---|
| WT Open | 75.346 | 25.285 | 0.722 |
| Omicron Open | 80.079 | 16.307 | 11.342 |
| WT Closed | 70.659 | 18.843 | 0.953 |
| Omicron Closed | 64.878 | 21.907 | 4.765 |
| RBD type | Glycan w/ RBD's RBM | Other Glycan w/ RBD's RBM | Glycan w/ remaining S1S |
| WT Open | 18.277 | 0.971 | 22.152 |
| Omicron Open | 12.207 | 0.928 | 10.157 |
| WT Closed | 10.061 | 14.105 | 42.516 |
| Omicron Closed | 14.867 | 21.938 | 21.409 |

Table S11: List of the replica used for the analysis with contour plots (Figures 4, 5, S7, S3, and S8), with the exceptions of H-bond analysis that considers all three replicas. Open-RBD trajectories are the same as those used in our previous work[2] in the presence of glycans.

| | closed-head | closed-RBM | Spike UDD |
|---|---|---|---|
| WT-PBL0 | 1 | 1 | 1 |
| Delta-PBL0 | 2 | 2 | - |
| Omicron-PBL0 | 1 | 3 | 2 |
| WT-PBL1 | 1 | 3 | 1 |
| Delta-PBL1 | 2 | 2 | - |
| Omicron-PBL1 | 3 | 2 | 2 |

### Supplementary Figures

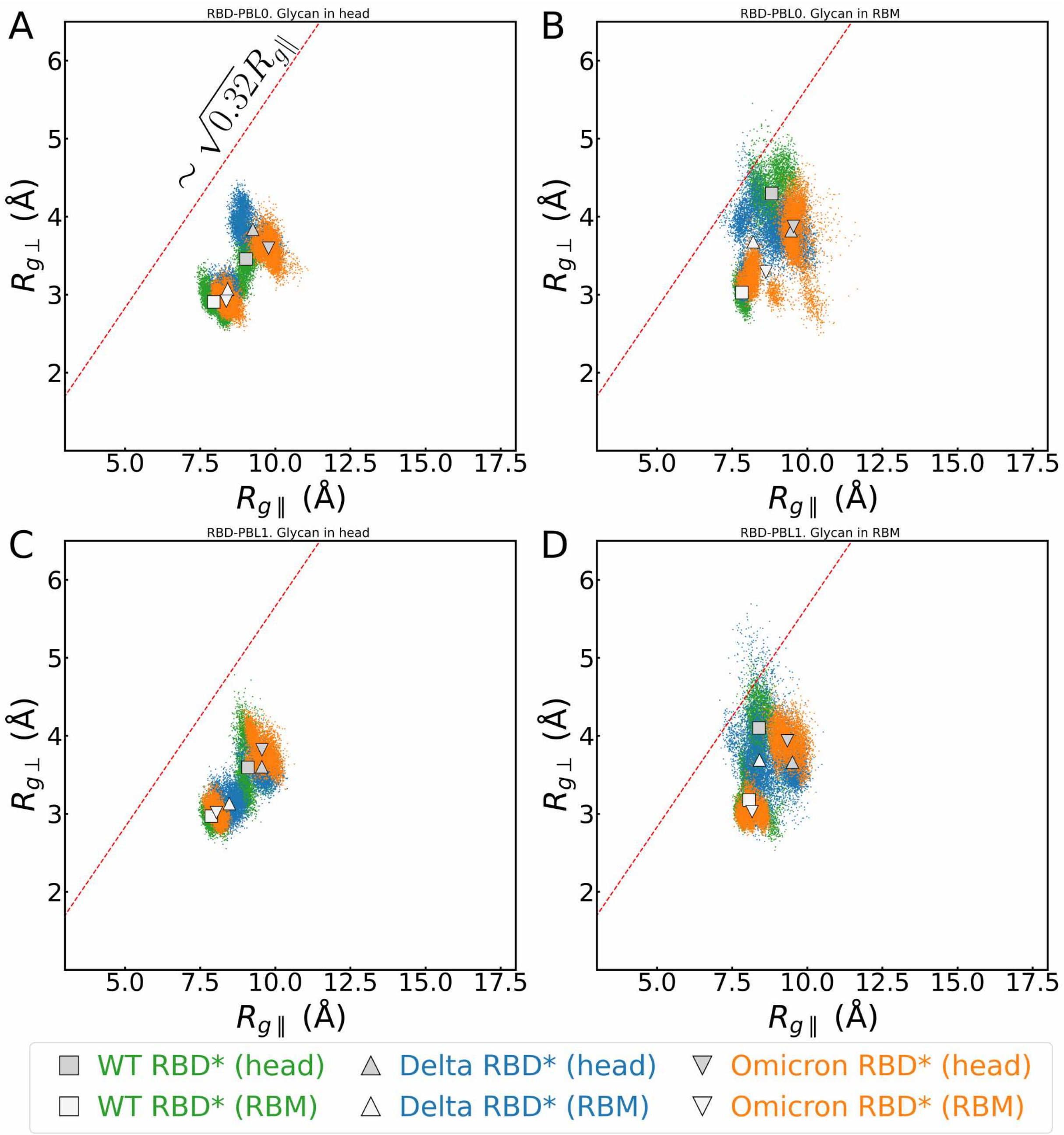


Figure S1: Parallel and perpendicular radii of gyration of the contact region of the head- and RBM-RBDs for WT (squares, green), Delta (triangles, orange), and Omicron (inverted triangles, blue) variants. Note that these are the contributions per region of white-filled colored symbols in Figure 2. The Red lines indicate the limit $\sim \sqrt{0.32} R_{g\|}$ for the semiflexible polymer regime. Panels show closed-RBD with the glycan between (A) a hydrophobic surface and the RBD's head, (B) a hydrophobic surface and the RBD's RBM, (C) a hydrophilic surface and the RBD's head, and (D) a hydrophilic surface and the RBD's RBM.

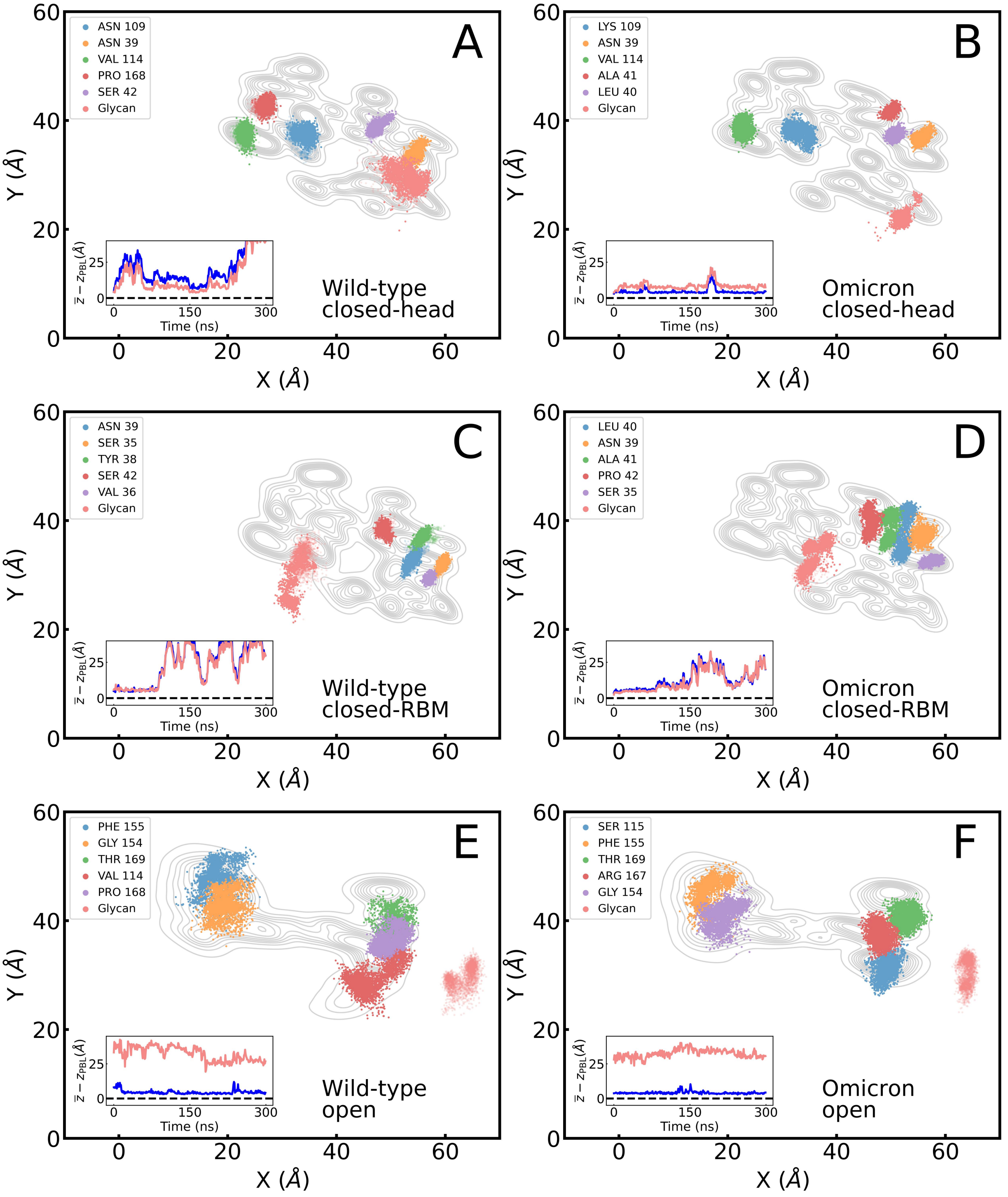


Figure S2: Footprint of the top five protein residues with the highest number of contacts to the hydrophilic surface and center-of-mass (COM) of the glycan for the closed- and open-RBD configurations for WT (left column) and Omicron variants (right column). Insets show the average distance of these residues (blue) and the glycan (red) to the surface. Panels display results for (A, B) closed-RBD with the glycan between the RBD head and the surface, (C, D) closed-RBD with the glycan between the RBM and the surface, and (E, F) open-RBD configuration. Contours of the contact regions of closed and open RBDs are shown as a reference map for the protein.

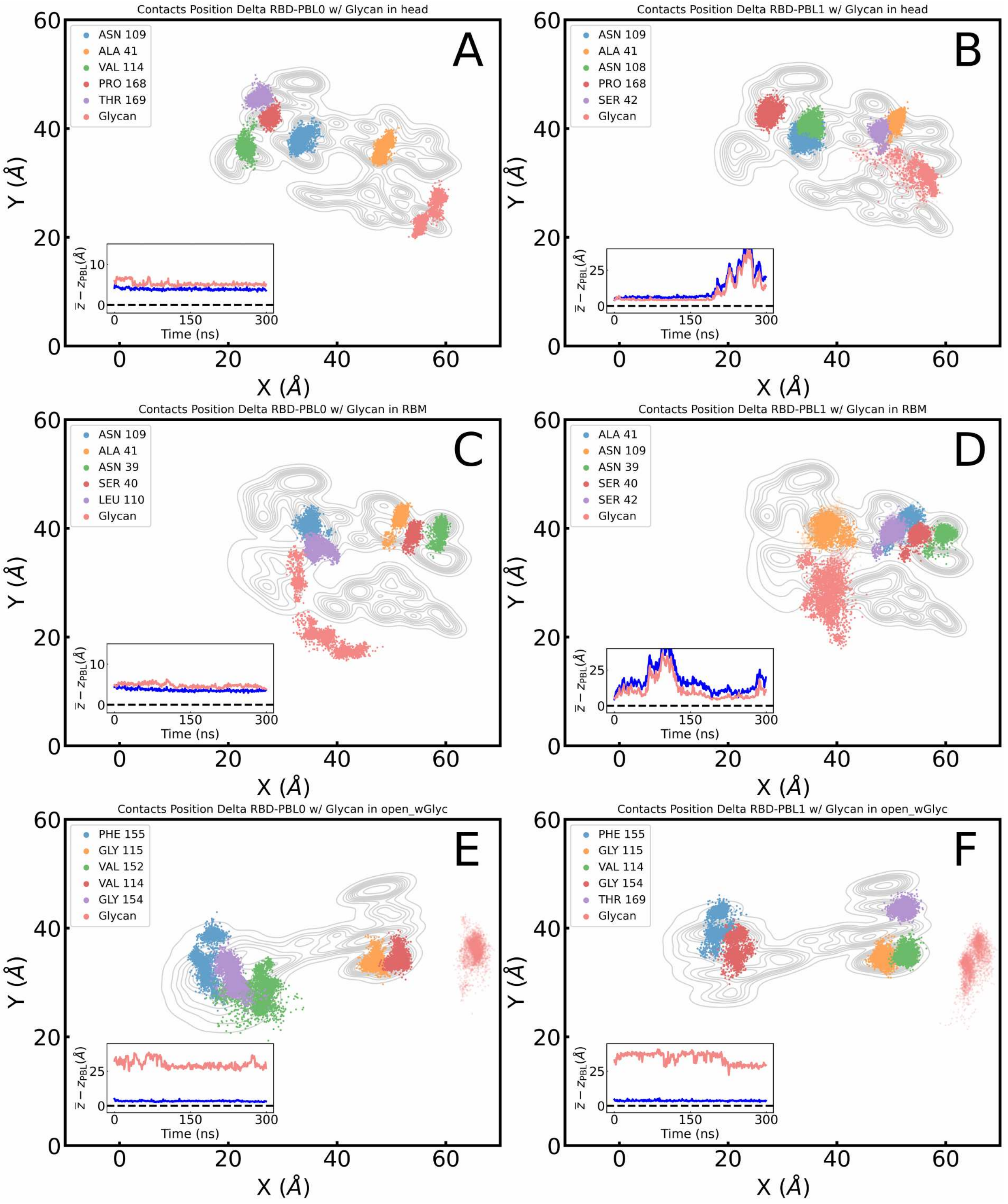


Figure S3: Footprint of the top five protein residues with the highest number of contacts to the hydrophobic (left column) and hydrophilic (right column) surfaces, and center-of-mass (COM) of the glycan for the closed- and open-RBD configurations for Delta variant. Insets show the average distance of these residues (blue) and the glycan (red) to the surface. Panels display results for (A, B) closed-RBD with the glycan between the RBD head and the surface, (C, D) closed-RBD with the glycan between the RBM and the surface, and (E, F) open-RBD configuration. Contours of the contact regions of closed and open RBDs are shown as a reference map for the protein.

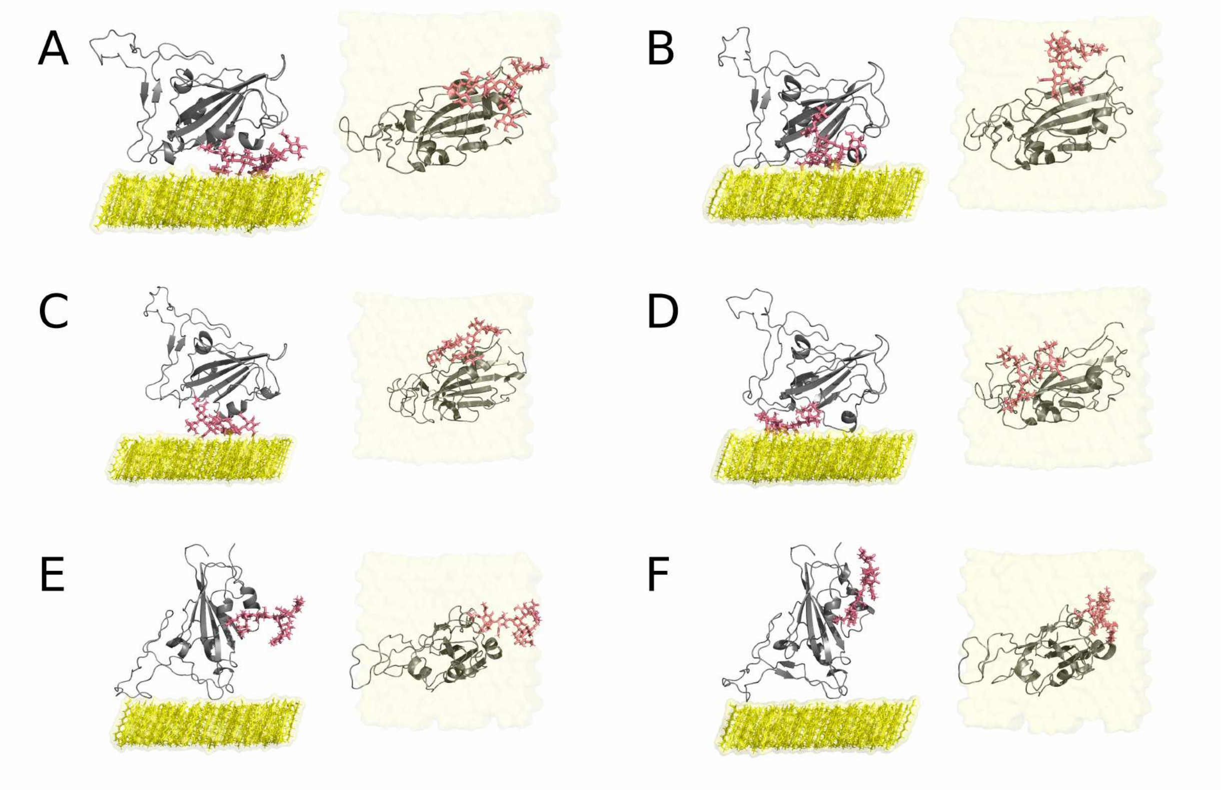


Figure S4: Snapshots of the RBDs with glycans onto the hydrophobic surface at 150ns, with the exception of (C) at 100ns, to evidence the anchoring of the protein via glycan. Replicas and panel ordering are consistent to simulations with the footprint contour plots (Figure 4 and 5.

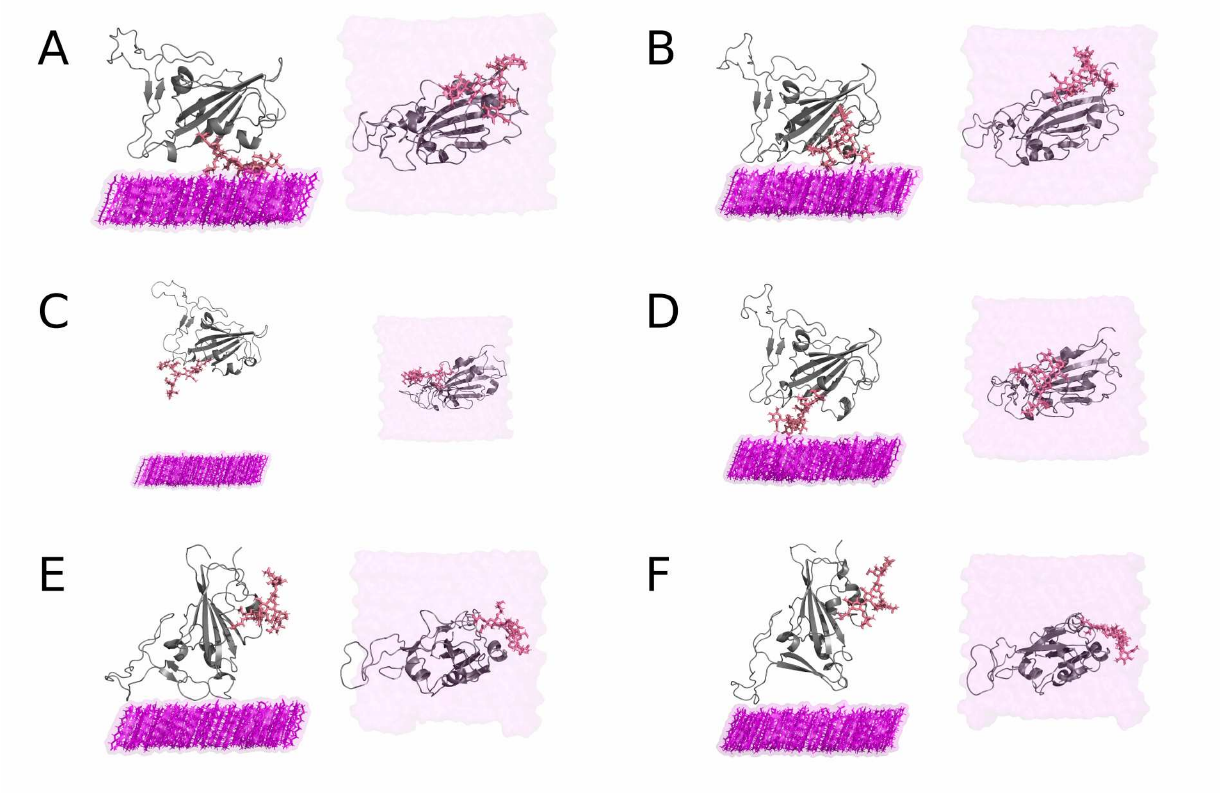


Figure S5: Snapshots of the RBDs with glycans onto the hydrophilic surface at 150ns. Replicas and panel ordering are consistent to simulations with the footprint contour plots (Figure S2 and S7).

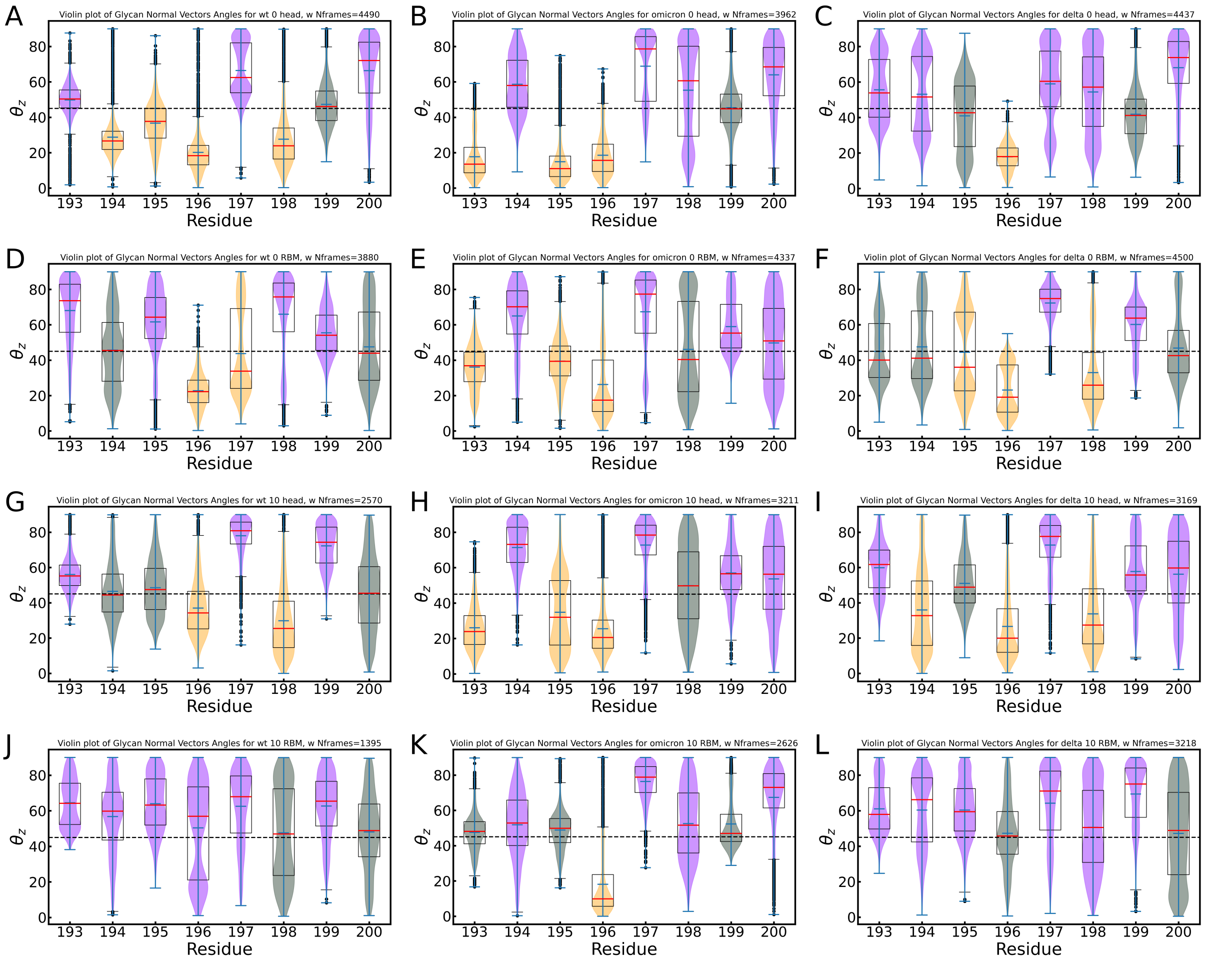


Figure S6: Violin and box plots of the angles ($\sigma$) of the sugar ring of the glycan with respect to the Z-axis, 0 is parallel and 90 is perpendicular to the Z-axis. Red lines of the box indicate the median value, and the blue bars of the violin plot indicate the max, min, and mean value. Colors of the violin plots highlight median values higher than 50 (purple, approximately perpendicular), lower than 40 (pastel orange, approximately parallel) or in the threshold between 40 and 50 degrees (grey). As a reference, a segmented line at 45 degrees is added. Black dots are data considered outliers by the box plots. Box and violin plots were computed using the Matplotlib functions. Panels (A-F) are from simulations onto the hydrophobic surface, and (G-L) onto the hydrophilic surface for WT(left), Omicron(center), and Delta (right) variants. (A-C, G-I) correspond to simulations with the glycan within the head-RBD and the surface, and (D-F, J-L) with the glycan within the RBM and the surface.

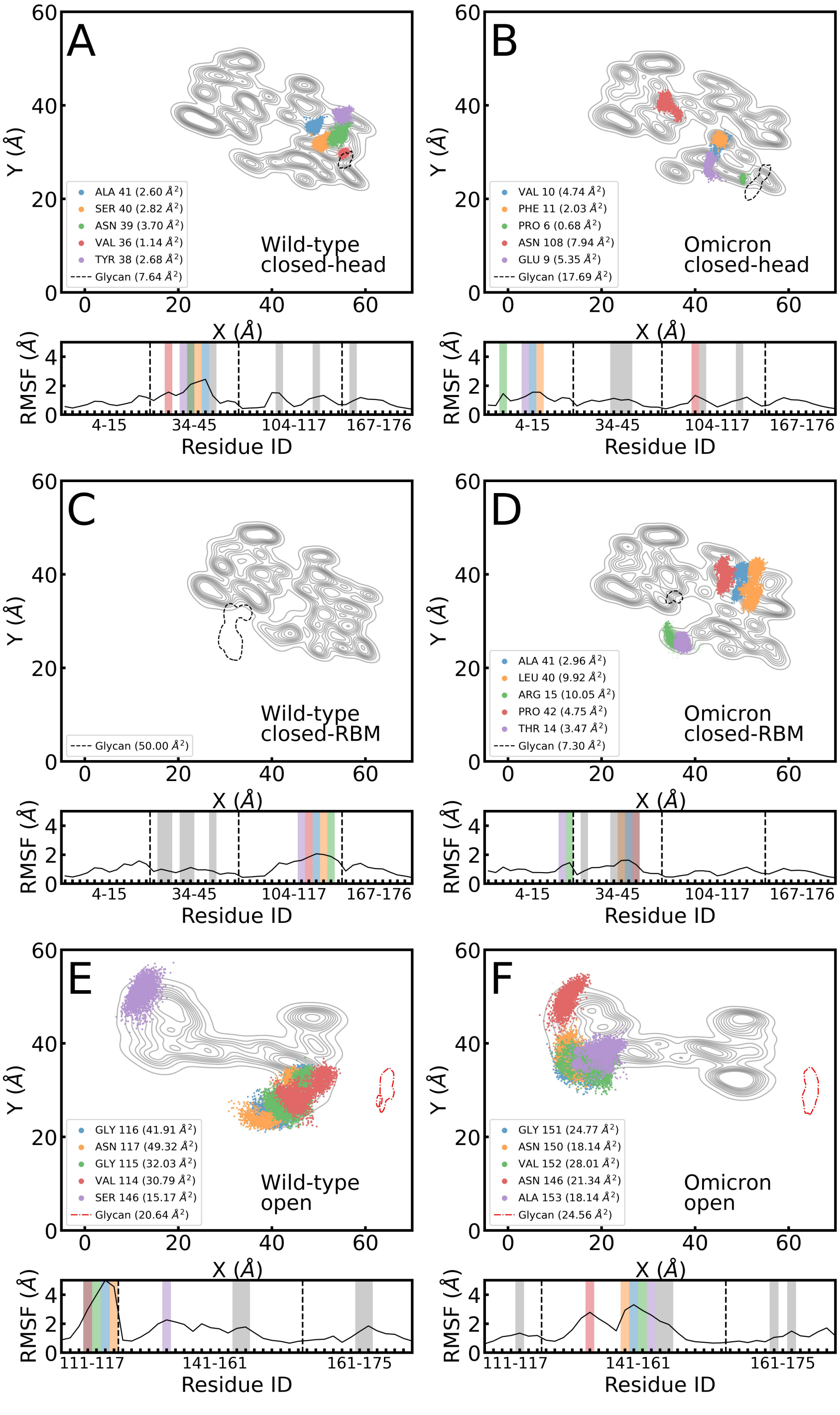

Figure S7: Footprint of the top five protein residues in the contact region with the highest RMSF values and area covered by glycan (dashed line) for the closed-RBD and open configurations onto the hydrophilic surface for WT (left column) and Omicron variants (right column). Panel ordering is consistent with Figures 5, 4, and S2. RMSF values for residues in the contact region are shown below each footprint, with colored regions highlighting the residues shown in the footprint and grey regions corresponding to residues with the highest number of surface contacts (Fig. S2). Red dot-dashed lines in (E) and (F) are a reminder that the glycan in the open conformations is not adsorbed (normal distance > 15 Å).

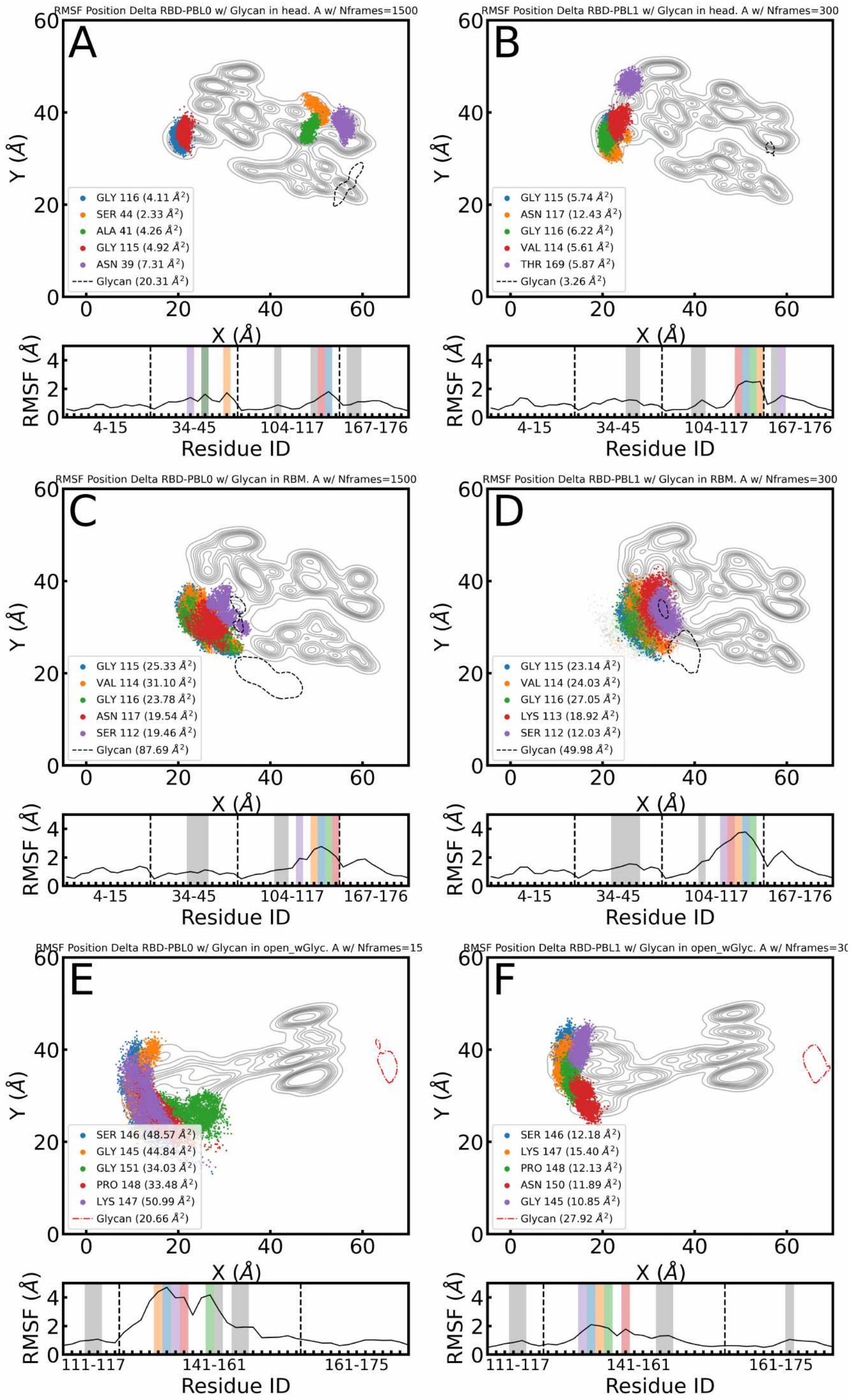


Figure S8: Footprint of the top five protein residues in the contact region with the highest RMSF values and area covered by glycan (dashed line) for the closed-RBD and open configurations the hydrophobic (left column) and hydrophilic (right column) surfaces for the closed- and open-RBD configurations for Delta variant. Panel ordering is consistent with Figure S3. RMSF values for residues in the contact region are shown below each footprint, with colored regions highlighting the residues shown in the footprint and grey regions corresponding to residues with the highest number of surface contacts (Fig. S3). Red dot-dashed lines in (E) and (F) are a reminder that the glycan in the open conformations is not adsorbed (normal distance > 15 Å).

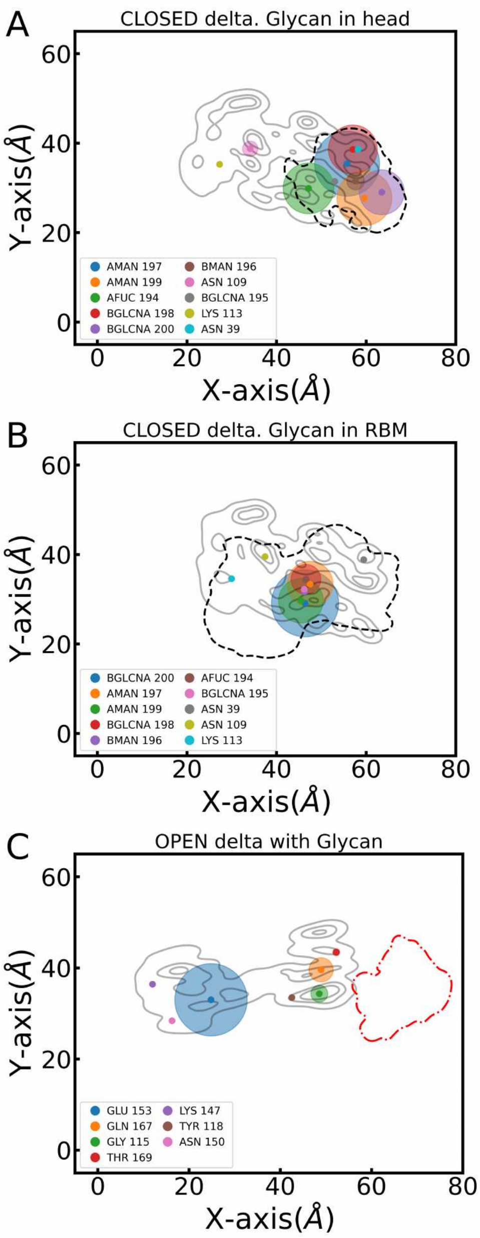


Figure S9: Hydrogen bonds occurrence between the protein and the surface during the simulation time (shown as %). H-bonds are represented in circles centered at the center-of-mass of each residue with proportional size to the percentage of H-bonding during the trajectory. The three contour levels in gray represent the mean positions of the protein, and the single contour in segment-black indicates the mean position of the glycan. Panels display results for (A) closed-RBD with the glycan between the RBD head and the surface, and (B) closed-RBD with the glycan between the RBM and the surface, and (C) open-RBD configuration.

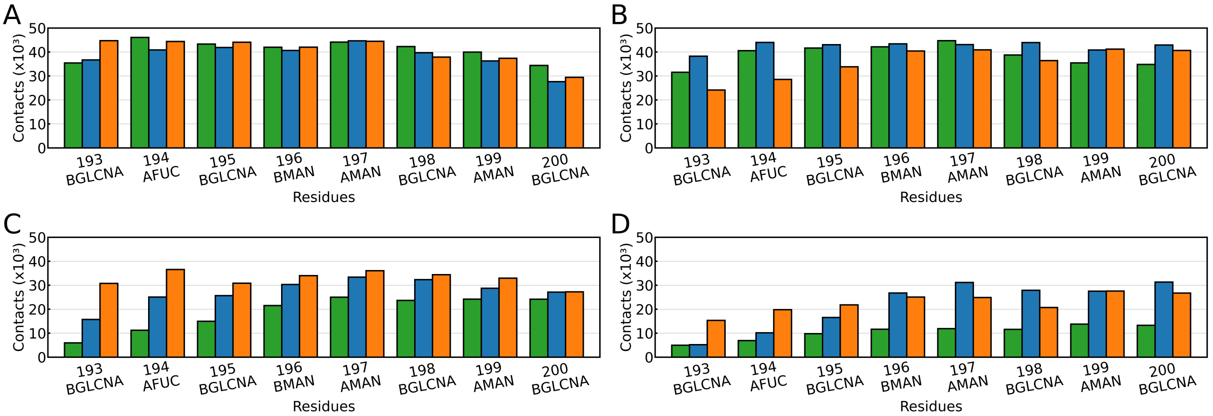


Figure S10: Glycan contacts with the surface when positioned between the surface and (A, C) the RBD head, or (B, D) the RBM region. Panels (A) and (B) correspond to a hydrophobic surface, while (C) and (D) correspond to a hydrophilic surface. Results are shown for the three VoCs; WT (green), delta (blue), and Omicron (orange).

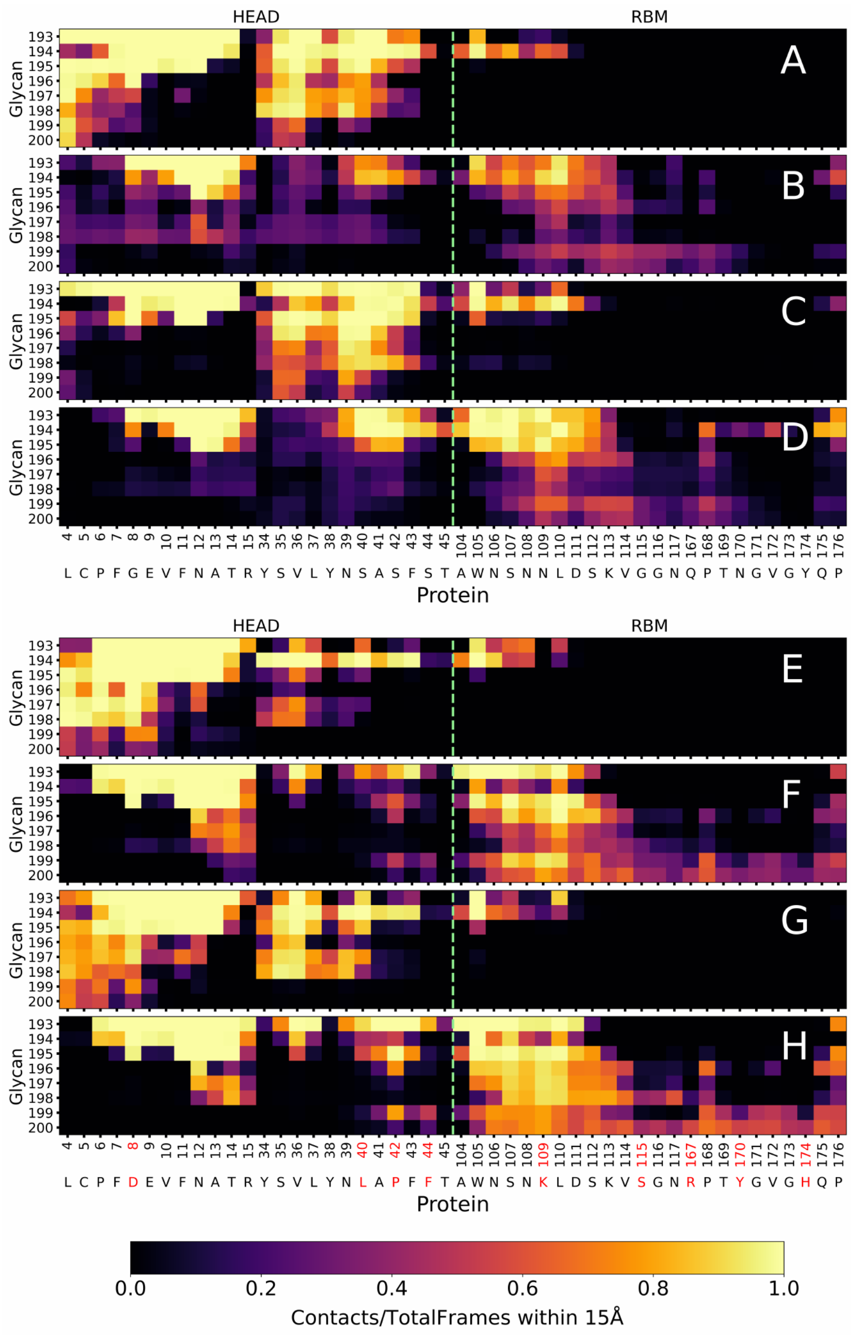


Figure S11: Contact maps between glycan and protein residues within the contact region for (A–D) WT and (E–H) Omicron. Panels (A, B, E, F) show results from simulations with a hydrophobic surface, while (C, D, G, H) correspond to a hydrophilic surface. In panels (A, C, E, G), the glycan is positioned between the RBD head and the surface; in (B, D, F, H), it is located between the receptor-binding motif (RBM) and the surface. A horizontal segmented lime green line in each map indicates the boundary separating residues in the RBD head from those in the RBM region of the RBD. Contacts are considered as residue-residue distances under 15Å.

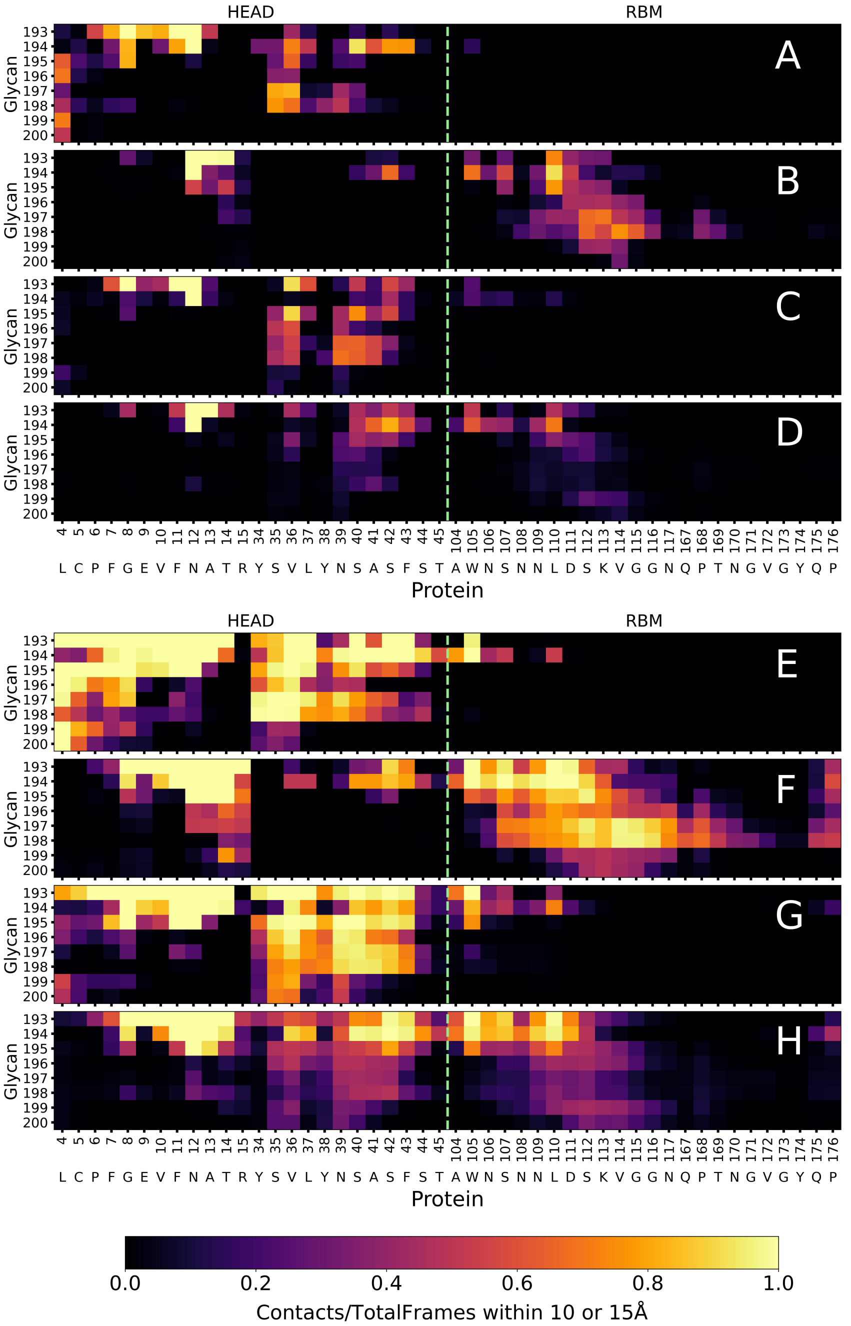


Figure S12: Contact maps between glycan and protein residues within the contact region for the Delta variant. A horizontal segmented lime green line in each map indicates the boundary separating residues in the RBD head from those in the RBM region of the RBD. Contacts are considered as residue-residue distances under (A-D) 10Å and (E-H) 15Å. Analogue to other figures with contact maps, panels (A, B, E, F) show results from simulations with a hydrophobic surface, while (C, D, G, H) correspond to a hydrophilic surface. In panels (A, C, E, G), the glycan is positioned between the RBD head and the surface; in (B, D, F, H), it is located between the receptor-binding motif (RBM) and the surface.

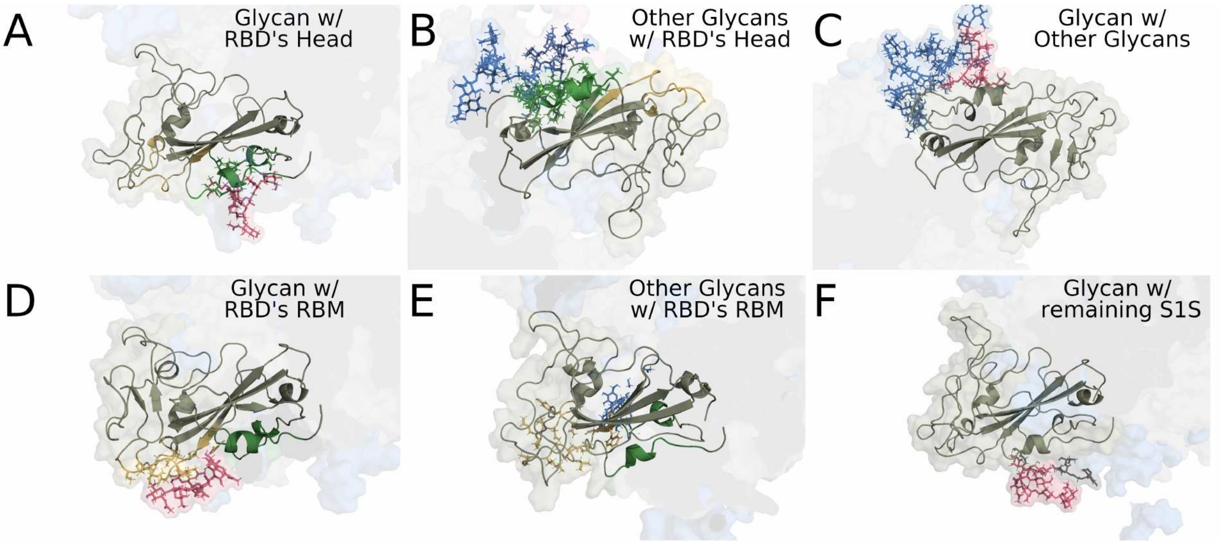


Figure S13: Snapshots showing the context of different regions of the RBD+glycan complex with the remaining S1 spike glycoprotein for open RBDs of WT. Glycan contacts with (A) the head and (D) the RBM of its RBD; other non-RBD glycans of the Spike with (B) the head and (E) the RBM; glycan with (C) other non-RBD glycans and with (F) protein residues of the remaining S1 Spike (S1S) glycoprotein. In the Figure, the RBD head is in green, the RBM region of the RBD is in orange, the glycan is in red, and the remaining S1 protein is in grey, and the remaining glycans of the S1 are in blue. Residues in sticks are at 7Å distance from the reference region. Reference regions : (A,B,D,E) Glycan of the RBDs, (C) head of RBD, and (F) RBM of the RBD.

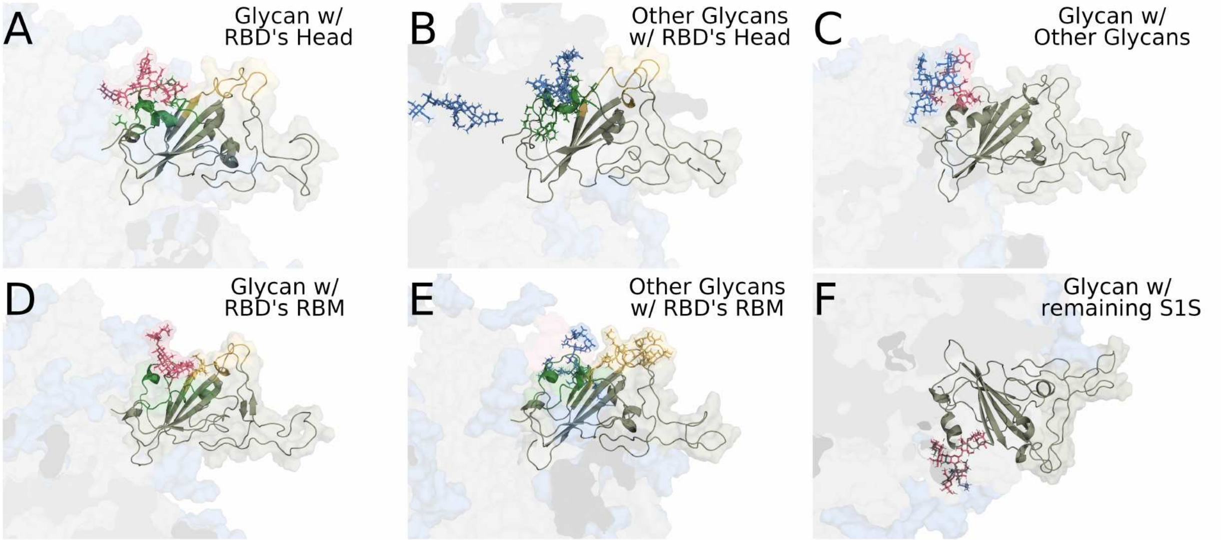


Figure S14: Snapshots showing the context of different regions of the RBD+glycan complex with the remaining S1 spike glycoprotein for open RBDs of Omicron. Panel display and color designation are the same as Figure S13.

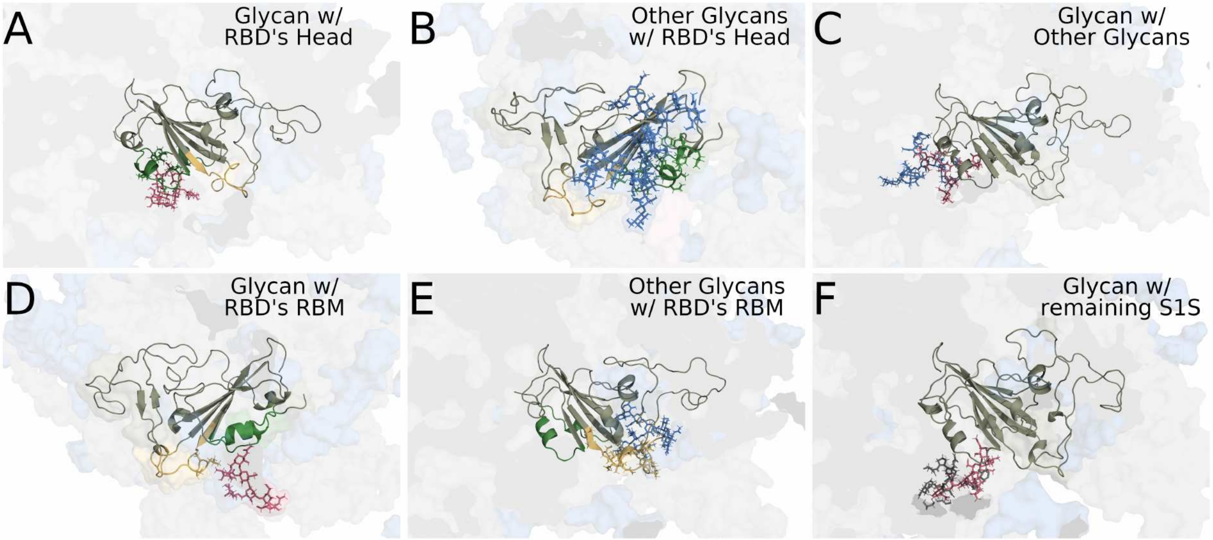


Figure S15: Snapshots showing the context of different regions of the RBD+glycan complex with the remaining S1 spike glycoprotein for closed RBDs of WT. Panel display and color designation are the same as Figures S13 and S14.

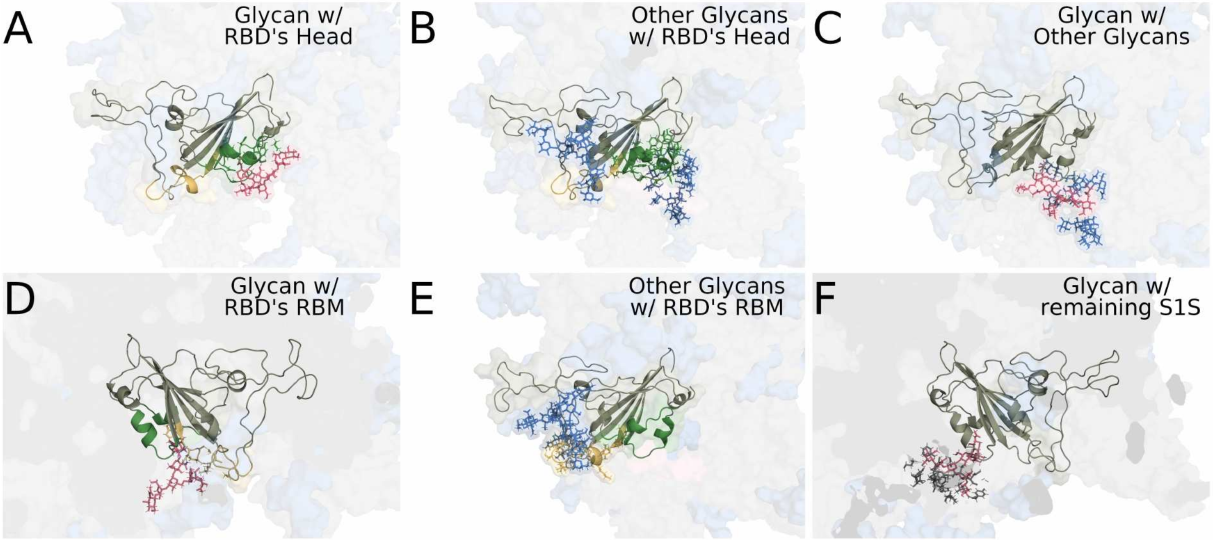


Figure S16: Snapshots showing the context of different regions of the RBD+glycan complex with the remaining S1 spike glycoprotein for closed RBDs of Omicron. Panel display and color designation are the same as Figures S13, S14, and S15.
